# Leukotrienes promote stem cell self-renewal and chemoresistance in acute myeloid leukemia

**DOI:** 10.1101/2022.02.07.479440

**Authors:** Alec W. Stranahan, Iryna Berezniuk, Sohini Chakraborty, Faye Feller, Mona Khalaj, Christopher Y. Park

**Author notes:** **Correspondence:** Christopher Y. Park, Department of Pathology, New York University Grossman School of Medicine, 522 First Avenue, New York, NY 10016. Equal contributions (Alec W. Stranahan, Iryna Berezniuk). Stanford University, Dept of Pathology and Institute for Stem Cell Biology and Regenerative Medicine (Mona Khalaj).

## Abstract

Acute myeloid leukemia (AML) is characterized by poor clinical outcomes due to high rates of relapse following standard-of-care induction chemotherapy. While many pathogenic drivers have been described in AML, our understanding of the molecular mechanisms mediating chemotherapy resistance remains poor. Therefore, we sought to identify resistance genes to induction therapy in AML and elucidated ALOX5 as a novel mediator of resistance to anthracycline based therapy. ALOX5 is transcriptionally upregulated in AML patient blasts in comparison to normal hematopoietic stem/progenitor cells (HSPCs) and ALOX5 mRNA, and protein expression is increased in response to induction therapy. *In vitro*, and *in vivo* genetic, and pharmacologic perturbation studies confirm that ALOX5 positively regulates the leukemogenic potential of AML LSCs, and its loss does not significantly affect the function of normal HSPCs. ALOX5 mediates resistance to daunorubicin (DNR) and promotes AML cell survival and maintenance through its leukotriene (LT) synthetic capacity, specifically via modulating synthesis of LTB4 and its binding to LTB receptor (BLTR). Our study reveals a previously unrecognized role of LTs in AML pathogenesis and chemoresistance, whereby inhibition of ALOX5 mediated LTB4 synthesis and function could be combined with standard chemotherapy, to enhance the overall therapeutic efficacy in AML.

**Key Points:**

- AML blasts overexpress and induce ALOX5 expression in response to daunorubicin
- 5-Lipoxygenase (ALOX5) promotes blast survival and LSC self-renewal.
- The effects of ALOX5 on AML blast survival and self-renewal are mediated by the LTB4-BLTR signaling axis.

## Introduction

AML results from the accumulation of genetic and epigenetic changes in HSPCs that lead to abrogation of their differentiation programs and accumulation in the bone marrow(1). AML is maintained by leukemia stem cells (LSCs) that possess the ability to self-renew as well as give rise to non-self-renewing blasts that comprise the vast majority of leukemia cells(2). LSCs have been shown to resist chemotherapy induced cell death, and form the cellular reservoir that mediates disease recurrence following therapy(2, 3). Thus, therapies designed with intent to cure AML should ideally eliminate LSCs without affecting normal hematopoietic cells including hematopoietic stem cells (HSCs).

Numerous studies have shown that AML blasts are metabolically distinct from normal HSPCs, exhibiting alterations in cellular stress, oxidative state, nutrient states, and responses to chemotherapy(4, 5). Indeed, blasts have been shown to exhibit alterations in glucose and amino acid metabolism. Recent studies in AML patients also have shown altered bone marrow(6) and plasma(7) lipid profiles and demonstrated that lipid metabolism likely contributes to progression, maintenance, and chemo-resistance(6, 8-10). While total serum fatty acids are decreased in AML patients, fatty acids such as arachidonic acid (AA) and its precursor, gamma-linolenic acid, are elevated. Moreover, elevated serum levels of AA and gamma-linoleic acid are associated with higher blast counts and poor overall survival in AML patients, suggesting a role of these fatty acids in AML blast growth and/or survival(7).

AA is metabolized through a series of enzymatic reactions that leads to the formation of bioactive lipid metabolites(11) through four enzymatic pathways: cyclooxygenase, lipoxygenase, cytochrome P450, and the anandamide pathways. One enzyme required for lipoxygenase synthesis, ALOX5, has been shown to promote chronic myeloid leukemia (CML) development and stem cell maintenance(12). In addition, ALOX5 dependent signaling promotes CML cell proliferation and viability(13, 14).

ALOX5 mediates an early step in the LT synthesis pathway, and thus is required for the synthesis of the entire family of LTs, which consists of LTA4, LTB4, LTC4, LTD4, and LTE4. Following conversion of AA to LTA4 by ALOX5, LTA4 is further processed by LTA4 hydrolase into bioactive LTB4; LTB4 is then converted into the cysteinyl LTs (LTC4, LTD4, and LTE4) through the actions of LTC synthase, transferase, and peptidase, respectively(15). LTs exert their biological functions by selectively binding to different receptors, with LTB4 binding to BLTR(16), and the cysteinyl LTs binding to CysLT1 and CysLT2(17). LTs have been shown to act as potent chemo-attractants that promote immune responses to pathogens, as well as a range of inflammatory and immune diseases(18-20). The pro-inflammatory activity of LTs and their correlation with tumor progression have been reported in multiple solid cancers, (21-23)(21-23)(21-23)with LTB4 contributing to cancer development and maintenance in a cell-extrinsic manner through excessive LT production by infiltrating leukocytes(21, 22). Despite these roles in solid cancers, the role of LTs in AML is poorly understood.

Out data indicate that ALOX5 mRNA expression is significantly increased in LSC-enriched and total blasts from AML patients. Given the role of the ALOX5/LT pathway in various human solid cancers and in CML stem cells(12, 24), we investigated the potential role of this pathway in AML pathogenesis. Genetic and pharmacologic loss-of-function experiments performed both *in vitro* and *in vivo* confirm that ALOX5 positively regulates the leukemogenic potential of LSCs. ALOX5 also mediates resistance to the anthracycline drug, daunorubicin (DNR)(25, 26), which is used as a component of conventional induction therapy. By inhibiting different LT receptors in AML cell lines and primary AML patient blasts, we show that ALOX5-LTB4 signaling, but not ALOX5-cysteinyl LT signaling, mediates ALOX5 dependent effects on AML blast survival in the presence and absence of DNR. Together, our studies establish the importance of the ALOX5-BLTR pathway in AML pathogenesis and chemo-resistance, and suggest that targeted inhibition of BLTR may be an important adjunct to standard induction therapy in AML.

## Methods

### In vitro cell line studies

MOLM-13 cells (DSMZ, cat no: ACC554) were transduced with lentiviruses expressing shRNAs against ALOX5 expressed using the pLKO-Puro backbone or empty vector as a control. Transduced clones were selected with puromycin. ALOX5 cDNA was ectopically expressed using the pLenti-Lox-GFP-mCherry vector backbone. GFP+ transduced clones were FACS-sorted and expression was induced by the addition of doxycycline (1.5 μg/ml) for 72 hours. Catalytically inactive ALOX5 was generated by making a single point mutation of HIS368SER in the ALOX5 cDNA construct as previously described (27, 28).

### ALOX5 inhibition and leukotriene receptor blocking studies

For cytotoxicity studies, cell death was assessed 72 hours after addition of DNR (Sigma, cat no: 30450) or Ara-C (LKT Laboratories, cat no: 147-94-4). For combination treatment studies with Zileuton (LKT Laboratories, cat no: Z3444, or Cayman Chemical, cat no: 10006967), cells were incubated with DNR (2, 3, 5, or 7nM, as indicated) and Zileuton (0-100μM) or DMSO (vehicle) for 3 days; Zileuton was replenished every 24 hours during the experiment.

For LT receptor blocking experiments, cells were treated with 4μM Montelukast, 1μM HAMI3379, or 5 μM LY293111 (Cayman Chemical, cat no: 10008318, 10580, and 10009768, respectively), or DMSO/MethylAcetate (vehicle) for the indicated time course. Cell death was assessed by flow cytometry using 0.1μg/mL propidium iodide (PI, Biolegend, cat no: 421301) and neutral beads (Beckman Coulter, cat no: B22804).

### *Alox5-/-* MLL-AF9 studies

Leukemic mice were generated by retroviral transduction of mouse HPSCs with the MLL-AF9 oncogene fusion as previously published(29). Double FACS-sorted Lin^-^Sca-1^+^c-Kit^+^ cells (LSK) were isolated from the bone marrow of either C57BL/6J WT or *Alox5-/-*. Following retroviral transduction with MLL-AF9, GFP+ cells were transplanted into irradiated primary C57BL/6J recipients as previously described(30). Leukemic mice were sacrificed, and bone marrow cells were isolated and utilized for secondary transplantation into C57BL/6J recipients.

For *in vitro* chemotherapy experiments, MLL-AF9+ blasts from the bone marrow were cultured in RPMI media supplemented with 10% FBS, 1% penicillin/streptomycin/L-glutamine (Gibco, cat no: 10378016), mIL-3 (10ng/mL), mSCF (20ng/mL), and mIL-6 (10ng/mL). Cells were passaged when >80% confluency was reached.

For *in vitro* DNR binding experiments, MLL-AF9+ blasts were treated with vehicle, 3 nM DNR, or 10 μM Zileuton for 72 hours followed by DNR-DNA intercalation assay described in the Flow Cytometry section.

Limiting dilution studies were performed by injecting increasing numbers of WT MLL-AF9 and *Alox5-/-* MLL-AF9 GFP+ bone marrow blasts into sublethally irradiated (475cGy) C57BL/6 mice via the retro-orbital venous sinus. Leukemia-initiating cell frequencies were determined using Extreme Limiting Dilution Analysis (ELDA) software(31).

## Results

### ALOX5 is upregulated in bulk and LSC-enriched AML blasts

Comparisons between AML blasts and normal HSPCs have identified numerous molecular pathways and biomarkers for disease detection, prognosis, and treatment(30, 32-34). We compared the transcriptome profiles of LSC-enriched (CD34+CD38-) and leukemia progenitor cell-enriched (LPC; CD34+CD38+) cell populations from AML patient bone marrow (BM) samples to normal human BM HSPC transcriptome profiles, using publicly available data in Gene Expression Commons(35, 36). Our analysis revealed that ALOX5 mRNA is expressed at significantly higher levels in both LSCs and bulk blasts compared to all HSPC populations (Figure 1A). Genes encoding key enzymes in other AA metabolic pathways, e.g. PTGS1/2 for cyclooxygenase pathway or CYP genes for the cytochrome P450 pathway were down-regulated in LSCs compared to HSPCs (https://gexc.riken.jp/; Supplemental Figure 1). Increased expression of ALOX5 mRNA in bulk blasts from different AML cytogenetic subtypes compared to normal HSPC population was confirmed in another publicly available dataset (http://servers.binf.ku.dk/bloodspot/; Figure 1B)(37). Evaluation of ALOX5 mRNA expression in functionally enriched CD99high LSCs from AML patient samples(38) revealed that CD99high LSC-enriched blasts overexpressed ALOX5 mRNA compared to CD99low non-LSCs by 1.9 fold (Figure 1C). Taken together, these data indicate that ALOX5 mRNA is overexpressed in LSC-enriched AML blasts and may contribute to LSC function.

**Figure 1.**
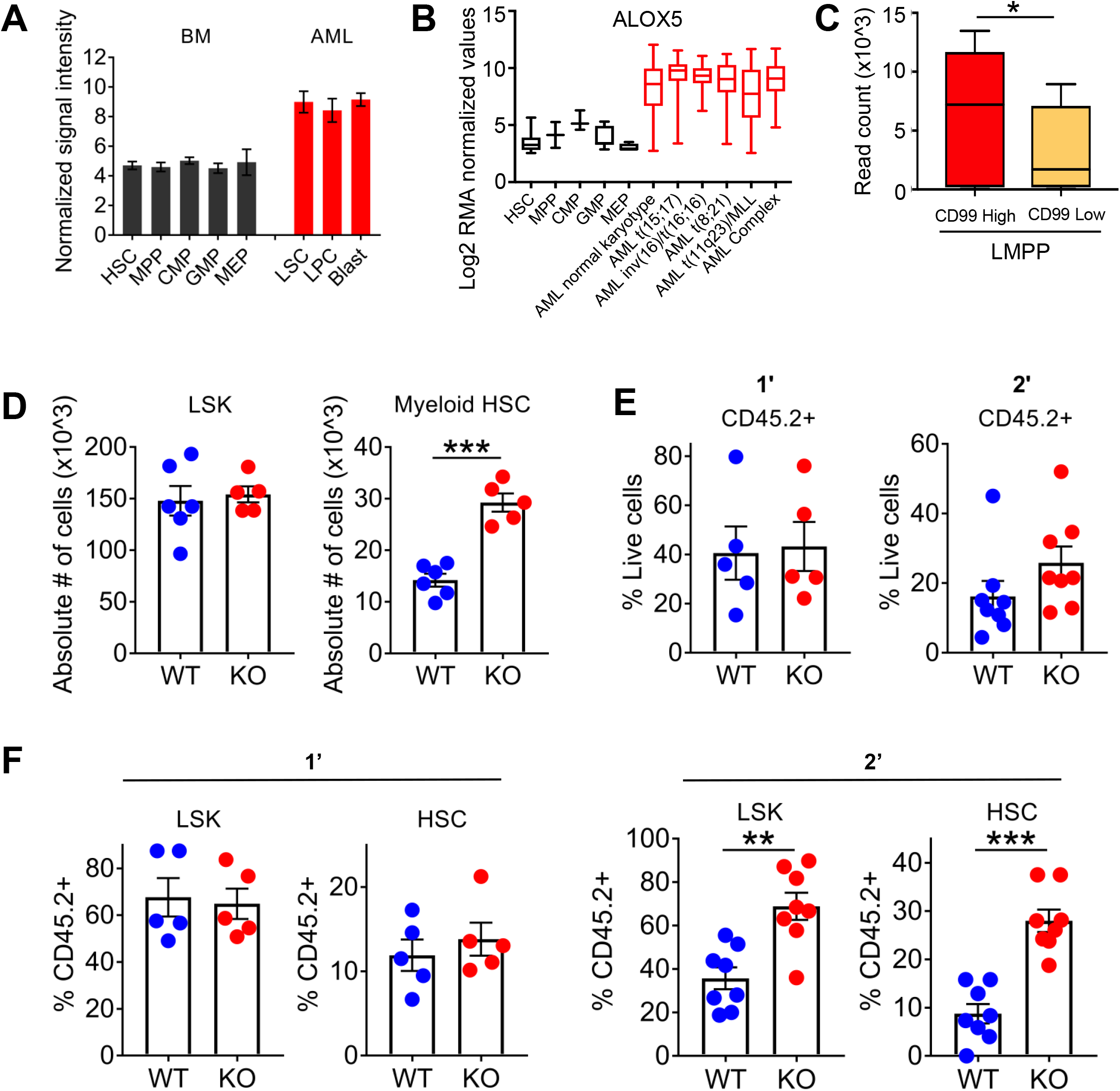
ALOX5 expression is upregulated in AML cells. (A) ALOX5 mRNA expression in normal HSPCs and AML cell types (from Gene Expression Commons, https://gexc.riken.jp). (B) ALOX5 mRNA expression in normal HSPCs and AML cell types (from http://servers.binf.ku.dk/bloodspot/). (C) ALOX5 mRNA expression in the LSC-enriched blasts (CD34+CD38−CD90−CD45RA+CD99+) and non-LSC blasts (CD34+CD38−CD90−CD45RA+CD99-). (D) Absolute cell numbers of LSK (Lin^-^Sca-1^+^c-Kit^+^) and myeloid-biased HSCs (Lin^-^Sca-1^+^c-Kit^+^CD34^-^CD150^high^) in the bone marrow of WT and *Alox5-/-* mice. (E) Flow cytometric analysis of recipient (CD45.1) and donor (CD45.2) cell chimerism in WT mice transplanted with WT or *Alox5-/-* HSCs, 16 weeks after transplantation. Bar graphs depict quantitation of mean CD45.2+ donor cells as a frequency of total propidium iodide-negative cells. (F) LSK frequency of total CD45.2+ cells and HSC frequency of total LSK cells in mice transplanted with WT and *Alox5-/-* donor cells. Primary (1’) and secondary (2’) transplantation. The results are presented as mean ± standard error mean (SEM). *P*-values were determined by the Student’s t-test: \**P* < 0.05, \*\**P* < 0.01, \*\*\**P* < 0.001.

### Loss of *Alox5* improves normal HSC function

Given the consistent high levels of expression of ALOX5 in AML, we assessed whether dysregulated ALOX5 expression alters normal hematopoiesis. ALOX5 mRNA expression was low in HSCs compared to more committed progenitors (http://servers.binf.ku.dk/bloodspot/; Figure 1B)(37). Evaluation of steady-state hematopoiesis showed that compared to WT controls, *Alox5-/-* mice showed a two-fold increase in the absolute number of immunophenotypically defined myeloid-biased HSCs (Lin-c-Kit+Sca-1+CD34-CD150high)(39) (Figure 1D). The absolute number of myeloid progenitor cells (Lin-c-Kit+Sca-1-) was reduced, largely due to a reduction in megakaryocyte-erythrocyte progenitors (MEP; Lin-c-Kit+Sca-1-CD34-CD16-) (Supplemental Figure 2A). There were no statistically significant differences in BM cell counts, lineage composition, or the frequency of other HSPC subsets in *Alox5-/-* mice (Supplemental Figure 2A-C), confirming observations from a previous study(40). In addition, peripheral blood (PB) leukocyte cell counts and lineage frequencies in *Alox5-/-* mice were comparable to WT controls, although *Alox5-/-* mice showed increased platelets and decreased red blood cells (Supplemental Figures 3A, 3B).

Although PB leukocyte numbers in *Alox5-/-* and WT mice were not different, the increase in CD150 high HSCs suggested an HSC myeloid lineage bias, and thus we tested HSC function and lineage output in long-term reconstitution experiments. While there was no significant difference in total CD45+ PB donor chimerism in primary recipients transplanted with WT and *Alox5-/-* HSCs, serial transplantation of c-Kit+ cells from primary recipients resulted in a trend towards a higher level of PB donor CD45+ chimerism in secondary recipients transplanted with *Alox5-/-* cells (Figure 1E). Mice transplanted with *Alox5-/-* donor cells also exhibited a two-fold higher frequency of donor-derived LSK cells and a three-fold higher frequency in donor HSCs compared to WT recipients (Figure 1F). Consistent with a myeloid lineage bias, secondary recipients receiving *Alox5-/-* cells exhibited a three-fold increase in GMPs (Lin-c-Kit+Sca-1-CD34+CD16+) and two-fold increases in PB monocyte and granulocyte chimerism at early time points following transplantation compared to WT recipients (Supplemental Figure 4A). No significant differences were observed in B- or T-cell chimerism in *Alox5-/-* and WT recipients (Supplemental Figure 4B). Together, these findings demonstrate that loss of ALOX5 promotes myeloid differentiation without negatively affecting HSC self-renewal.

### ALOX5 promotes LSC self-renewal

To test the potential role of ALOX5 in leukemogenic transformation, we tested the ability of MLL-AF9 to transform WT and *Alox5-/-* HSPCs using leukemia colony formation assays. MLL-AF9+ HSPCs exhibited significantly lower clonogenic capacity and produced fewer cells compared to WT counterparts (Figure 2A).

**Figure 2.**
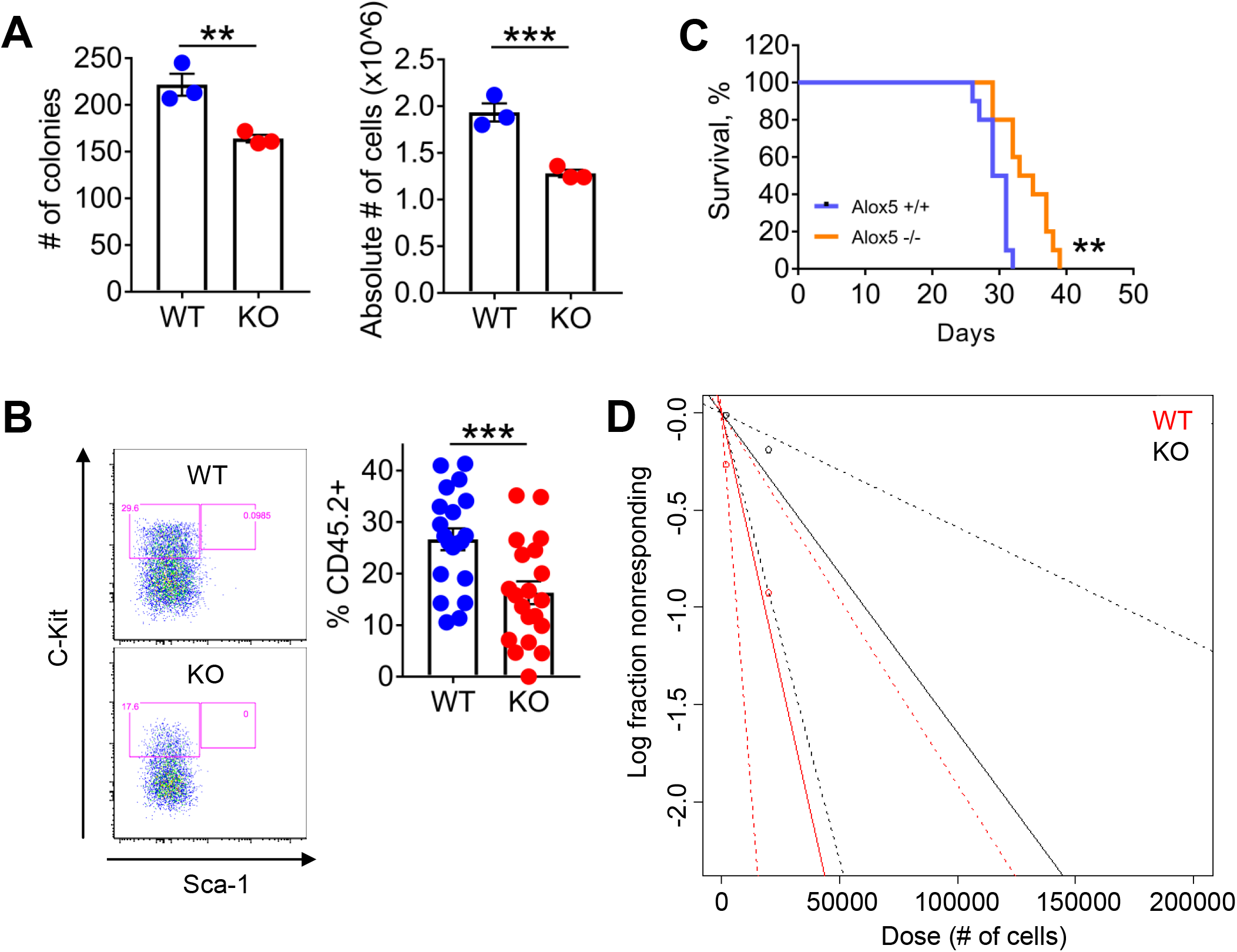
Loss of ALOX5 affects LSC self-renewal and maintenance. (A) Number of colonies (left panel) and absolute number of cells (right panel) as determined by CFU assay after plating WT or *Alox5-/-* MLL-AF9 blasts. (B) Representative dot plots of bone marrow GFP+ cells from mice engrafted with WT and *Alox5-/-* MLL-AF9 leukemia (left panel), frequencies of engrafted L-GMP cells (GFP^+^Lin^-^Sca-1^-^c-Kit^+^CD34^+^CD16^+^) (right panel). (C) Kaplan-Meier curves of mice engrafted with WT or *Alox5-/-* MLL-AF9 blasts (n=10/group). (D) LIC frequency as determined by limiting dilution assay. The results are presented as mean ± SEM. *P*-values were determined by the Student’s t-test: \*\**P* < 0.01, \*\*\**P* < 0.001.

After secondary plating, *Alox5-/-* MLL-AF9+ transformed blasts essentially failed to form colonies, suggesting that ALOX5 functions to promote LSC self-renewal (Supplemental Figure 4C). To test this possibility *in vivo*, we transplanted mouse recipients with equal numbers of WT or *Alox5-/-* HSPCs transduced with MLL-AF9 retrovirus. Mice transplanted with *Alox5-/-* blasts exhibited reduced frequencies of LSCs (L-GMP; Lin-c-Kit+Sca-1-CD34+CD16+)(41) compared to those transplanted with WT blasts (Figure 2B). Moreover, mice engrafted with *Alox5-/-* AMLs exhibit a significant extension in survival compared to mice transplanted with WT blasts (Figure 2C). Limiting dilution transplantation assays revealed that *Alox5-/-* blasts exhibited a decreased leukemia-initiating cell (LIC) frequency of 1/60,780 compared to 1/18,409 in MLL-AF9 WT AML (Figure 2D). Collectively, these results demonstrate that ALOX5 promotes LSC self-renewal.

### *Alox5-/-* blasts exhibit altered transcriptional programs

To investigate the molecular mechanisms by which ALOX5 promotes leukemogenesis and LSC maintenance, we performed RNA-sequencing on MLL-AF9+ WT and *Alox5-/-* leukemic (GFP+) cells. *Alox5-/-* blasts showed globally altered transcriptional changes, with *Alox5-/-* blasts showing 635 differentially expressed genes (DEGs) (fold change ≥ 2, p.adj ≥ 0.05) (Figure 3A). Gene Set Enrichment Analysis (GSEA) of the DEGs revealed enrichment of signaling pathways required for LSC survival and MLL-AF9+ blast maintenance in WT vs *Alox5-/-* blasts, including an MLL associated gene signature, oxidative phosphorylation, DNA repair, cell cycle checkpoints, PI3K/AKT/MTOR signaling, and targets of Myc, p53, and MDM4 (Figure 3B). In contrast, *Alox5 -/-* blasts were enriched for apoptotic response and inflammatory response gene signatures (Figure 3C). These gene expression changes support ALOX5’s role in promoting multiple critical pathways required for AML LSC self-renewal and growth.

**Figure 3.**
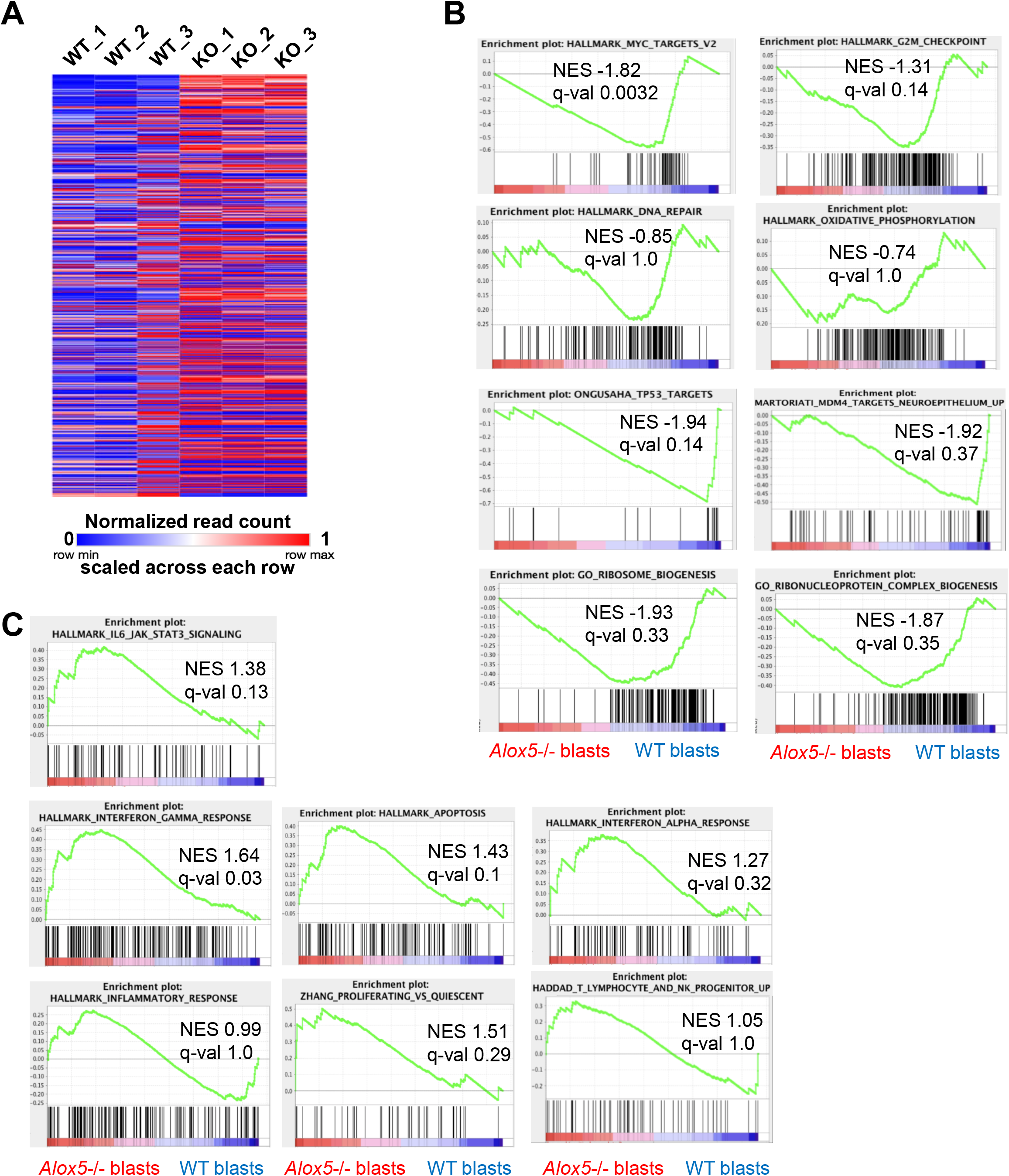
ALOX5 depletion affects signaling pathways required for LSC survival and function. (A) Heatmap of differentially expressed genes between WT and *Alox5-/-* MLL-AF9 leukemic blasts (n=3). (B) Enrichment plots of GSEA pathways upregulated (positive enrichment score) in WT vs *Alox5-/-* leukemic cells. (C) Enrichment plots of GSEA pathways upregulated (positive enrichment score) in *Alox5-/-* MLL-AF9+ blasts compared to WT MLL-AF9+ leukemic cells.

### ALOX5 promotes chemoresistance and is induced in response to DNR

Given ALOX5’s role in promoting AML LSC self-renewal, we next investigated if ALOX5 deficiency may sensitize blasts to chemotherapy. shRNA mediated knockdown of ALOX5 in MOLM-13 cells increased their sensitivity to DNR, but not AraC or the two drugs in combination (Figure 4A). To confirm the role of ALOX5 in promoting resistance to DNR, we first treated WT and *Alox5-/-* MLL-AF9+ blasts with DNR. Consistent with the role of ALOX5 in promoting blast survival, *Alox5-/-* MLL-AF9+ blasts showed a 28% reduction in cell viability compared to WT blasts; this decrease in cell viability was even more pronounced following treatment with DNR-treated blasts with a 74% decrease in live *Alox5-/-* MLL-AF9+ cells compared to WT blasts (Figure 4B). Overexpression of ALOX5 in MOLM-13 cells nearly abrogated DNR-induced cell death, confirming that ALOX5 is sufficient to mediate chemoresistance (Figure 4C). To confirm that the ALOX5 knockdown phenotypes were not due to off-target effects, we ectopically expressed ALOX5 in ALOX5 knockdown cells. These studies confirmed that ALOX5 knockdown inhibits leukemic cell growth in the absence of DNR and that ectopic ALOX5 cDNA expression is sufficient to rescue the growth defect following ALOX5 knockdown (Figure 4D) as well as reverse the chemo-sensitizing effect of ALOX5 loss (Figure 4E). Consistent with the chemo-protective function of ALOX5, DNR sensitivity was not rescued by ectopic ALOX5 expression when ALOX5 was knocked down more efficiently (Figures 4E, 4F), indicating that a minimal level of ALOX5 expression is essential for leukemia cell survival in response to DNR. Finally, DNR treatment of MOLM-13 cells induced ALOX5 mRNA expression (Figure 4G). Thus, ALOX5 is not only overexpressed in AML blasts, but is induced in order to mediate its chemoresistance and cell survival phenotypes in response to DNR treatment.

**Figure 4.**
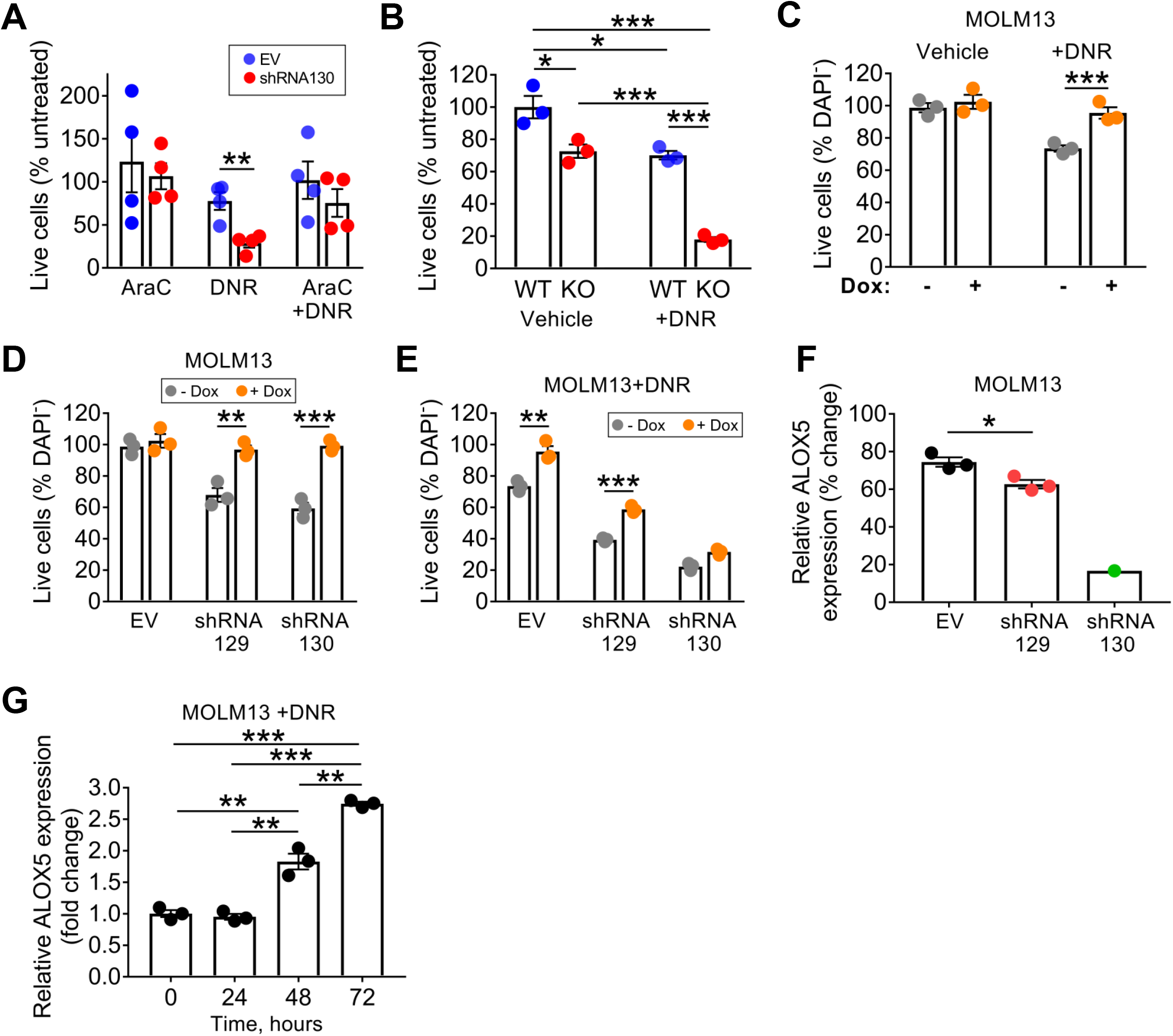
ALOX5 is induced with DNR treatment and ectopic ALOX5 expression increases chemoresistance. (A) Survival of MOLM-13 cells expressing either empty vector or ALOX5 shRNA following treatment with 300 nM AraC, 7 nM DNR, or both, for 72 hours. (B) Live cell frequency of WT and KO MLL-AF9+ blasts following treatment with 7 nM DNR for 72 hrs. (C) Survival of MOLM-13 cells ectopically expressing doxycycline (Dox)-inducible ALOX5 following 72 hrs treatment with DNR. (D) Live cell frequency following ectopic ALOX5 expression of two independent ALOX5 shRNAs (129, 130). (E) Live cell frequency of MOLM-13 cells expressing ALOX5 shRNAs (129, 130) complemented with ectopically expressed ALOX5 in the presence of DNR. (F) Normalized ALOX5 mRNA expression following ALOX5 shRNA mediated knock-down as measured by qPCR. (G) Time course of ALOX5 mRNA expression following exposure to DNR. Live cells were measured as % of DAPI-negative cells (%DAPI-), normalized to untreated cells. The results are presented as mean ± SEM. *P*-values were determined by the Student’s t-test (compared to empty vector (EV) control (A), no Dox (B-D) or time 0 (F): \**P* < 0.05, \*\**P* < 0.01, \*\*\**P* < 0.001.

To test whether DNR resistance requires ALOX5’s enzymatic activity, we expressed a catalytically-inactive mutant form of ALOX5 in myeloid leukemia cell lines(27, 28). While expression of wildtype ALOX5 resulted in decreased cell death in response to DNR (Figure 5A), mutant ALOX5 did not, despite the fact that WT and mutant transcripts and proteins were expressed at comparable levels (Figure 5B). In addition, pharmacologic inhibition of ALOX5 using the FDA-approved inhibitor Zileuton resulted in a significant increase in cell death in the presence of DNR (Figure 5C). Finally, we exogenously administered LTs (combination of LTB4, LTC4, LTD4, LTE4) to MOLM-13 cells treated with DNR, which rescued the chemo-sensitizing effects of ALOX5 knockdown and Zileuton treatment (Figures 5D, 5E). Sensitivity to DNR correlated with the amount of DNR DNA intercalation, since DNR intercalation increased slightly upon pharmacologic inhibition of ALOX5 with Zileuton in MLL-AF9+ WT blasts, and even more in ALOX5-deficient *Alox5-/-* MA9+ blasts (Figure 5F). The increase in DNR DNA binding did not appear to be due to differences in drug transporter activity, as no significant alteration in DNR efflux was observed between WT and *Alox5-/-* leukemic MLL-AF9+ blasts over a wide range of DNR concentrations (Supplemental Figure 5A), and the expression of canonical drug transporters in AML including Abcb1b (P-gp), Abcc1 (MRP1), and Abcg2, and other genes associated with drug resistance were not increased in *Alox5-/-* versus WT blasts (Supplemental Figure 5B). Together, these studies demonstrate that ALOX5 is required for chemoresistance in leukemic blasts.

**Figure 5.**
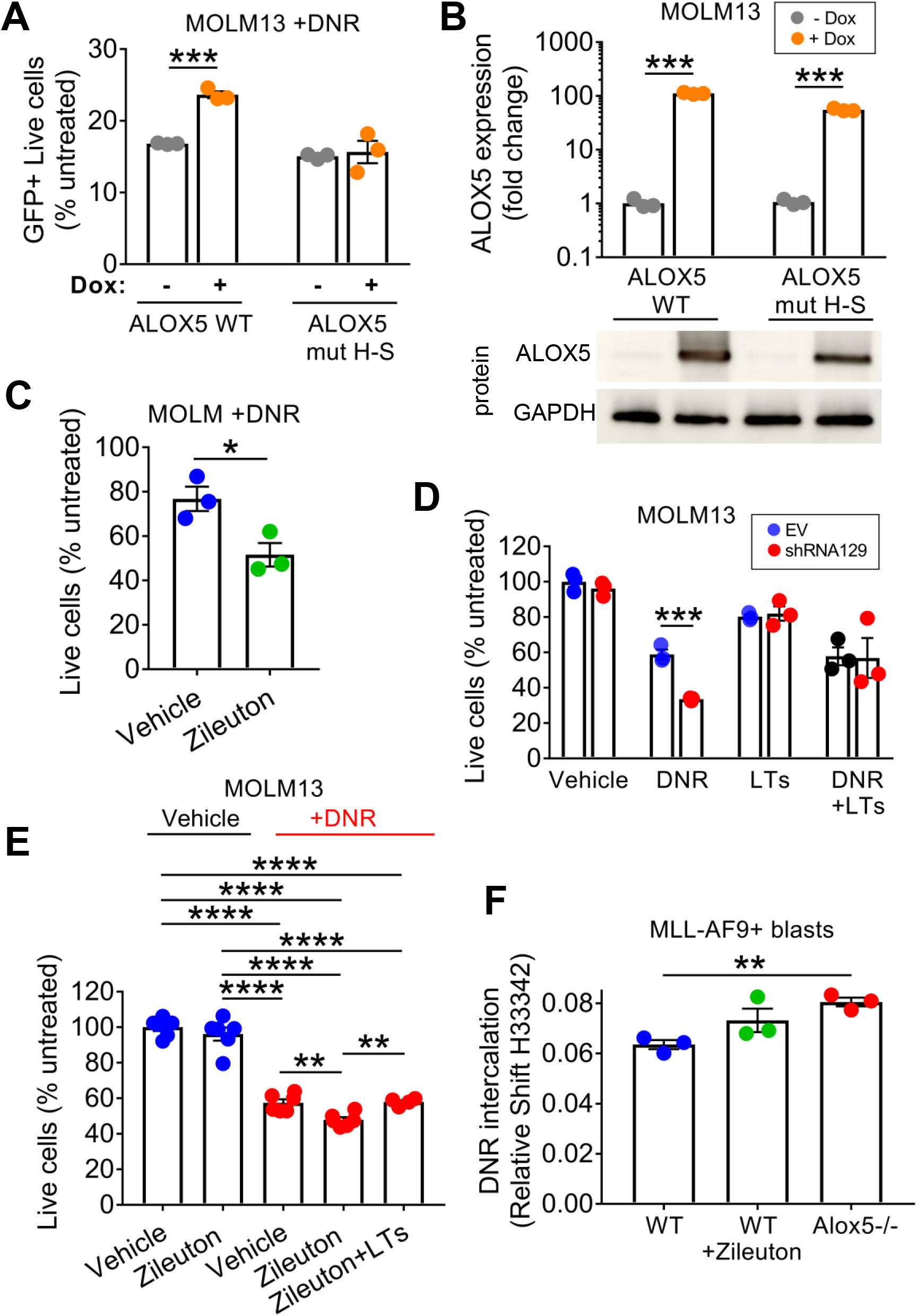
The enzymatic activity of ALOX5 is required for chemoresistance of AML cells. (A) Survival of MOLM-13 cells expressing inducible WT or mutant ALOX5 following treatment with DNR. (B) WT or mutant ALOX5 expression in MOLM-13 cells as measured by qPCR (top panel) and western blot (bottom panel). (C) Survival of MOLM-13 cells following treatment with either 1 μM Zileuton or DMSO vehicle in the presence of 3nM DNR for 72 hrs. (D) Survival of MOLM-13 cells containing scrambled (scr) or ALOX5 shRNA (shRNA 129) vectors after treatment with DNR, a mixture of LTs (LTB4, LTC4, LTD4, and LTE4), or DNR and LTs. (E) Survival of MOLM-13 cells after treatment with either 1 μM Zileuton or DMSO vehicle in the presence of 3nM DNR in combination with 1 μM Zileuton and 25 nM LTs. (F) DNR DNA binding was assessed by measuring the spectral shift of Hoechst 33342 peak emission in WT and *Alox5-/-* MLL-AF9 leukemic cells treated with Zileuton *in vitro*. Live cells were measured as %DAPI- or %PI-cells, normalized to untreated cells. The results are presented as mean ± SEM. *P*-values were determined by the Student’s t-test (compared to no Dox (A, B), vehicle (C), sc shRNA (D), and WT (F): \*\**P* < 0.01, \*\*\**P* < 0.001.

### LTB4-BLTR axis mediates the ALOX5 dependent phenotypes

ALOX5 generates multiple LT family members that exert their biological effects by binding to three unique receptors: LTC4, LTD4, and LTE4 bind to CysLT1 and CysLT2, while LTB4 binds to BLTR(42, 43). To determine which LTs may mediate ALOX5 dependent chemoresistance, we first evaluated the expression of the different LT receptors. Analysis of gene expression data from TCGA AML, and BEAT AML cohorts revealed that BLTR mRNA expression is 3-fold higher than CysLT1, and CysLT2 (Figure 6A, Supplemental Figures 6A, 6B). BLTR mRNA expression is significantly correlated with the expression of ALOX5 (Supplemental Figures 6A, 6B, 6C), and LTA4H, the gene encoding LTA4 hydrolase required for the synthesis of LTB4 from LTA4 (Supplemental Figure 6C). In contrast, expression of LTC4 synthase (LTC4S), which is required for the synthesis of LTC4 from LTA4, did not correlate with ALOX5 expression. DNR treatment of MOLM-13 cells induced surface BLTR expression, but did not alter the expression of the other LT receptors (Figure 6B). DNR treatment also induced surface BLTR expression in WT MLL-AF9+ blasts, but not in *Alox5-/-* blasts (Supplemental Figure 7A). Together, these data indicate that multiple components of the ALOX5-LTB4-BLTR signaling axis are coordinately regulated at the transcriptional level in AML, both at steady-state and in response to DNR.

**Figure 6.**
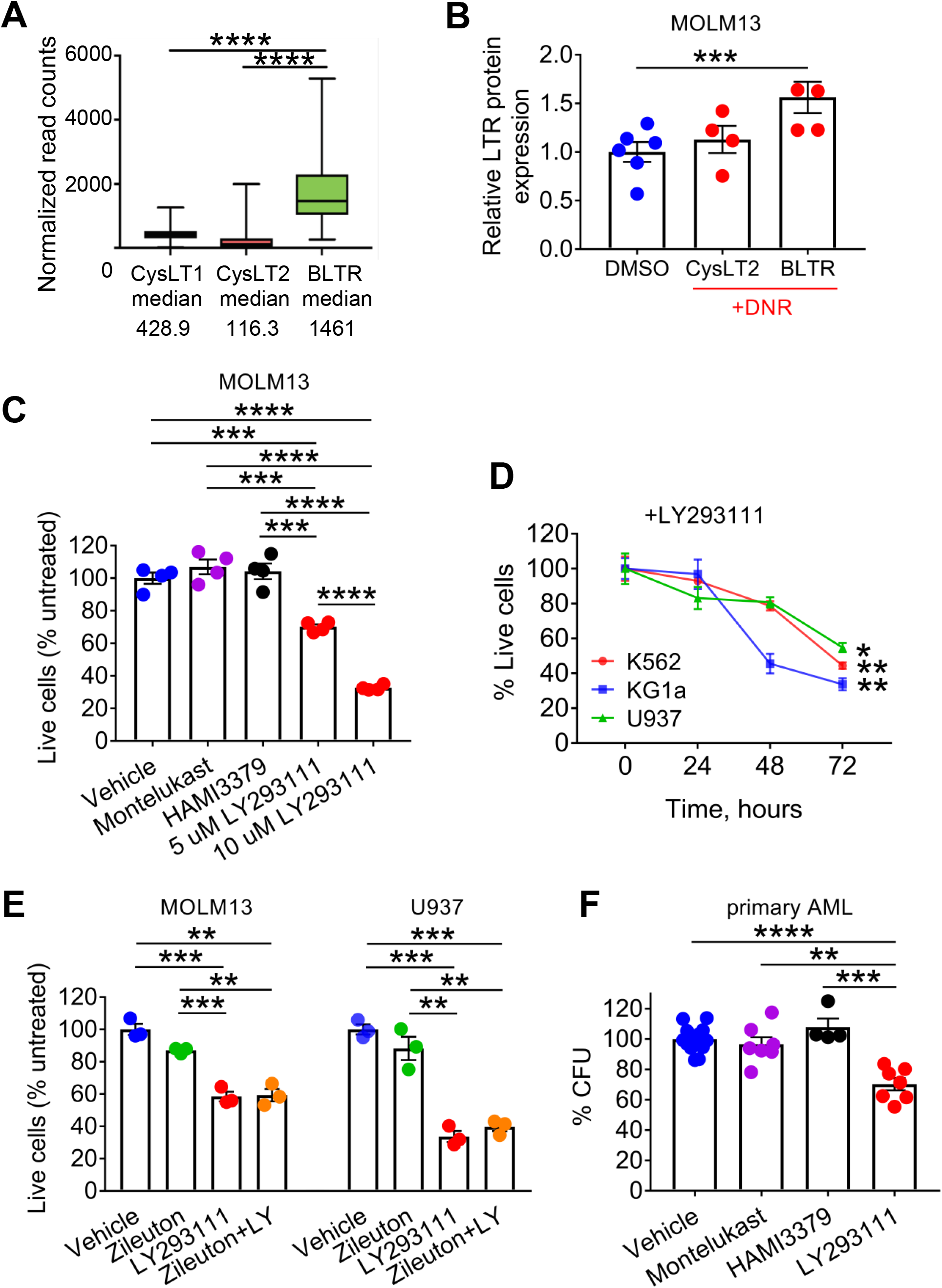
BLTR axis downstream of ALOX5 is necessary for AML cell survival. (A) Expression of LT receptor transcripts in AML patient samples in the TCGA cohort. (B) Normalized expression of LT receptor proteins after treatment of MOLM-13 cells with 3 nM DNR for 24 hrs, as measured by flow cytometry. Fold change expressed relative to vehicle-treated cells. The level of CysLT1 was comparable to unstained control after treatment. (C) Survival of MOLM-13 cells after treatment with CysLT1 receptor antagonist Montelukast (4 μM), CysLT2 receptor antagonist HAMI3379 (1 μM), or BLTR antagonist LY293111 (5 or 10 μM). (D) Time course of survival of K562, KG1a, and U937 cells after treatment with 5 μM LY293111. (E) Survival of MOLM-13 and U937 cells after treatment with LY293111, Zileuton, or both, for 72 hours. Live cells were measured as %PI-cells, normalized to untreated cells. (F) Number of colonies after plating primary AML blasts in Methocult containing vehicle, Montelukast, HAMI3379, or LY293111 (n=6 independent patient samples). The results are presented as mean ± SEM. *P*-values were determined by the Student’s t-test: \**P* < 0.05, \*\*\**P* < 0.001, \*\*\*\**P* < 0.0001.

To test the potential to pharmacologically target the ALOX5-LTB4-BLTR signaling for therapeutic benefit, we inhibited BLTR function with LY293111, a second-generation selective BLTR inhibitor(44).

Treatment with LY293111 induced cell death in AML and CML blast crisis cell lines (Figures 6C, 6D) as well as in primary AML patient blasts, even in the absence of DNR (Supplemental Figure 7B). However, neither inhibition of CysLT1 with the potent antagonist Montelukast(45), nor inhibition of CysLT2 with the selective inhibitor HAMI3379(46), affected AML cell or blast survival (Figures 6C, Supplemental Figure 7B). Blocking BLTR signaling with LY293111 exerted a greater effect on AML cell survival than inhibition of ALOX5 with Zileuton, and combined inhibition with Zileuton and LY293111 did not further increase cytotoxicity (Figure 6E). In leukemic colony formation assays, BLTR inhibition with LY293111 significantly decreased the clonogenicity of primary AML patient blasts, whereas CysLT1 and CysLT2 receptor antagonists did not (Figure 6F). Further supporting the role of LTB4-BLTR signaling in leukemogenesis, exogenously administered LTB4 increased the clonogenic capacity and colony size of WT and *Alox5-/-* MLL-AF9+ leukemic blasts compared to untreated counterparts (Supplemental Figures 7C, 7D). Confirming the role of LTB4 in promoting resistance to DNR, we treated cells with LY293111 and found that it sensitized AML cell lines to DNR cytotoxicity (Supplemental Figures 7E, 7F). Recapitulating the effect of ALOX5 knockdown or enzymatic inhibition, inhibition of BLTR signaling by LY293111 also induced increased DNR binding in MOLM-13 cells (Supplemental Figure 7G) as well as in primary AML patient blasts (Supplemental Figure 7H). Collectively, these data indicate that the effects of ALOX5 on LSC self-renewal and resistance to DNR treatment is largely dependent on LTB4-BLTR signaling.

## Discussion

We have identified the ALOX5-LTB4-BLTR axis as a novel pathway required for leukemic transformation, LSC self-renewal, and therapeutic responses to anthracycline therapy in AML. ALOX5 expression is significantly increased in human AML blasts compared to normal HSPCs, including blasts functionally enriched in LSCs activity, and is induced in response to DNR, a mainstay of standard induction chemotherapy. Additionally, loss-of-function studies using genetic or pharmacologic inhibition approaches demonstrate the critical role of ALOX5 in maintaining the leukemogenic potential of MLL-AF9 transduced HSPCs, as well as promoting survival in transformed blasts both *in vitro* and *in vivo. Alox5-/-*. Since ALOX5 is not required for normal hematopoiesis, and loss of ALOX5 even enhances normal HSC self-renewal, targeting the ALOX5-LTB4-BLTR axis in combination with anthracycline treatment is a novel and excellent candidate therapeutic strategy in AML.

Previous studies have demonstrated a role for LTs in CML, as leukocytes from CML patients produced more cysteinyl LTs compared to healthy donors(47), and selective CysLT1 antagonists induced dose-dependent inhibition of CML cell line growth(13). Moreover, it was shown that depletion of ALOX5 affected leukemic stem cell self-renewal in a mouse model of CML(12). However, these studies did not assess the role of LTs in AML, and they did not investigate which LTs regulate CML cell growth. In addition, BLTR blockade did not consistently inhibit cell growth in CML cell lines. LT function has been investigated in AML, but these studies neither evaluated effects of LT on LSC self-renewal, nor did they evaluate the functional role of specific LTs in AML blasts or confirm the role of ALOX5 through overexpression of catalytically inactive form or using knockout approaches. Moreover, these studies only considered specific genetic subtypes of AML such as APL and t(11q23)/MLL leukemia(48-53), and they even suggested that catalytically active ALOX5 is not required for growth promoting effects in APL. In contrast, our studies indicate that ALOX5 promotes blast survival and self-renewal at steady state as well as in response to chemotherapy, but is dispensable for normal HSC self-renewal. Moreover, our studies show that the catalytic activity of ALOX5 is required to mediate AML cell chemoresistance phenotypes and we show that among the LT synthetic products of ALOX5, LTB4 most prominently contributes to AML blast survival and growth, both in the absence or presence of DNR. In addition, the effects of ALOX5 on these phenotypes likely are not AML genotype specific, as transcriptional upregulation of the major regulatory components of the ALOX5-LTB4-BLTR pathway are increased in human AML regardless of cytogenetic subtype, including within LSC-enriched fractions. Indeed, in addition to demonstrating the role of the ALOX5-LTB4-BLTR pathway in mediated resistance to DNR in the MA9+ mouse model of AML, we also demonstrated similar dependencies on the ALOX5-LTB4-BLTR pathway using patient AML blasts representing non-MLL rearranged AMLs (Figures 6B, 6E, 7E).

The effects of the ALOX5-LTB4-BLTR pathway on promoting blast survival in response to chemotherapy are, at least in part, mediated by the ability of the ALOX5-LTB4-BLTR pathway to inhibit DNA binding by DNR. While the exact mechanism by which LTB4 mediates DNR DNA binding is not presently clear, it does not appear to be due to alterations in drug import or export. Indeed, pharmacologic blockade of BLTR, but not CysLT1 or CysLT2, inhibited the proliferation and clonogenic capacity of all AML cell lines and primary cells tested following DNR treatment (Figures 6C-E, 7A). Given that the BLTR antagonist LY293111 (Etalocib)(54), an FDA-approved drug that has been shown to reduce cancer cell survival and growth in multiple tumor models(55-57), can significantly inhibit blast survival at doses that are achievable *in vivo* without promoting systemic toxicity(58) suggests that a therapeutic strategy combining LY293111 with anthracycline-containing therapy may improve therapeutic efficacy.

Given the robust loss-of-function leukemogenic phenotypes observed in the *Alox5-/-* background including decreased leukemic proliferation, increased cell death, and increased sensitivity to DNR, we were somewhat surprised to find that inhibiting BLTR signaling with the LY293111 had a greater effect on AML cell survival than inhibition of ALOX5 with Zileuton and that the combination of both drugs did not exhibit an additive effect towards AML cell killing. This is likely explained by the robust activity of LY293111 in inhibiting LTB4-BLTR interactions as well as the lack of complete inhibition of ALOX5 enzymatic activity and potential off-target effects following Zileuton treatment, which has been reported in CML and pancreatic cancer cell lines, even when using clinically relevant concentrations of Zileuton(13, 59).

In summary, our studies strongly suggest that LTs play a prominent role in the development of AML, LSC self-renewal, and blast responses to chemotherapy, specifically through LTB4 finding to BLTR. Given that multiple studies suggest that AML patients will benefit from therapeutic strategies that combine drugs targeting novel therapeutic targets with standard chemotherapy(20, 60), we propose the ALOX5-LTB4-BLTR pathway represents an attractive target for treatment of newly diagnosed AML, either singly or in combination with anthracycline-containing therapies.

## Acknowledgments

The authors thank Drs. Neil Rosen and Ross L. Levine for scientific discussions and Memorial Sloan Kettering Cancer Center for use of its Shared Services (Flow Cytometry Core, Integrated Genomics Operation, and Molecular Cytology Core).

## Authorship Contributions

Contribution: A.W.S., F.F., I.B., and C.Y. P. conceived and designed the study; A.W.S. and M.K. acquired the mouse data; A.W.S. and C.Y.P. analyzed and interpreted the mouse data; I.B. and S.C. carried out BLTR related experiments and analysis; S.C. performed formal analysis of the transcriptomic data; I.B. performed and analyzed the experiments involving primary AML specimens; A.W.S., I.B., and C.Y.P. wrote, reviewed, and/or revised the manuscript; M.K. provided technical or material support; and C.Y.P. supervised the study.

## Figure Legends

**Figure S1.**
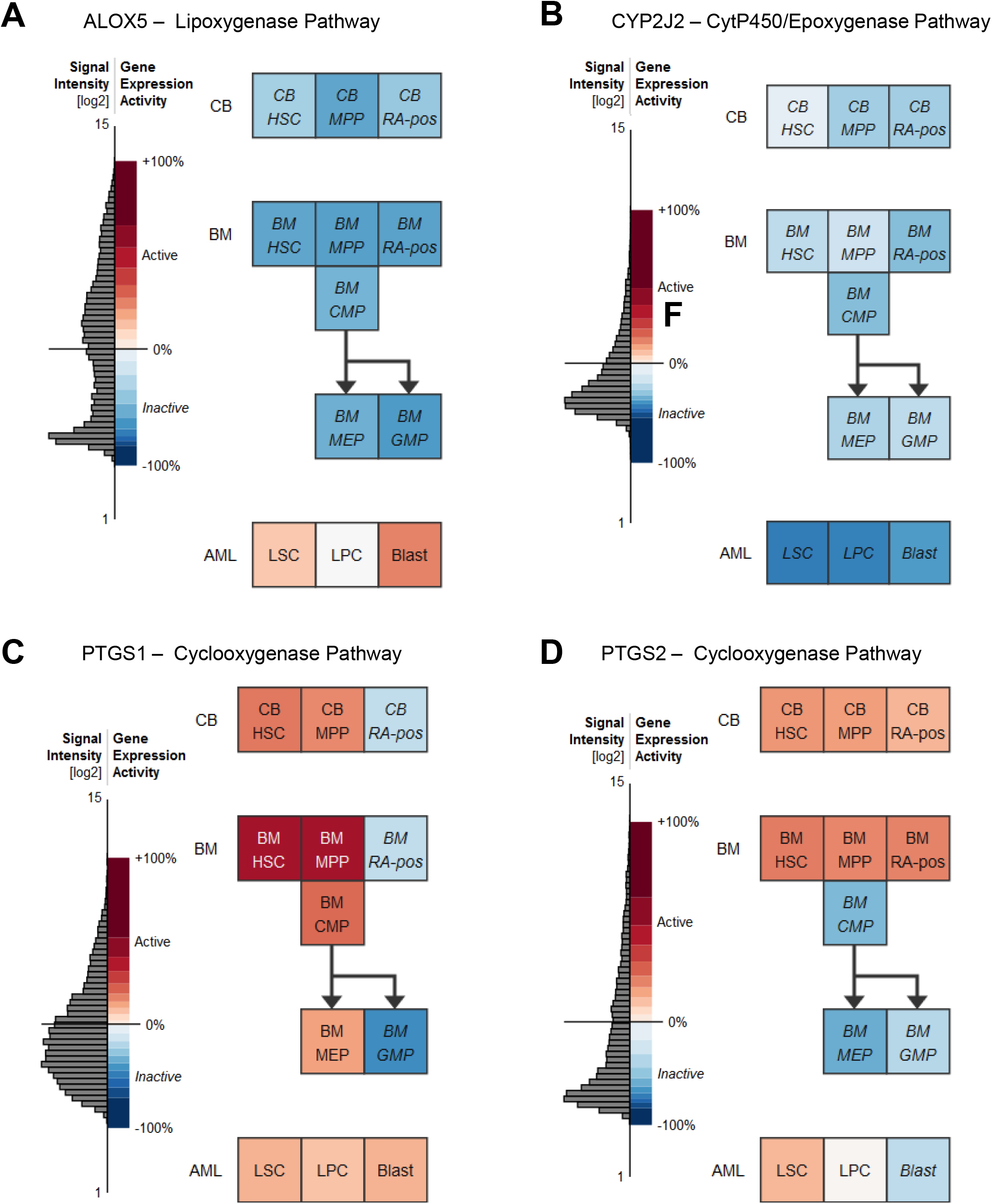

**Figure S2.**
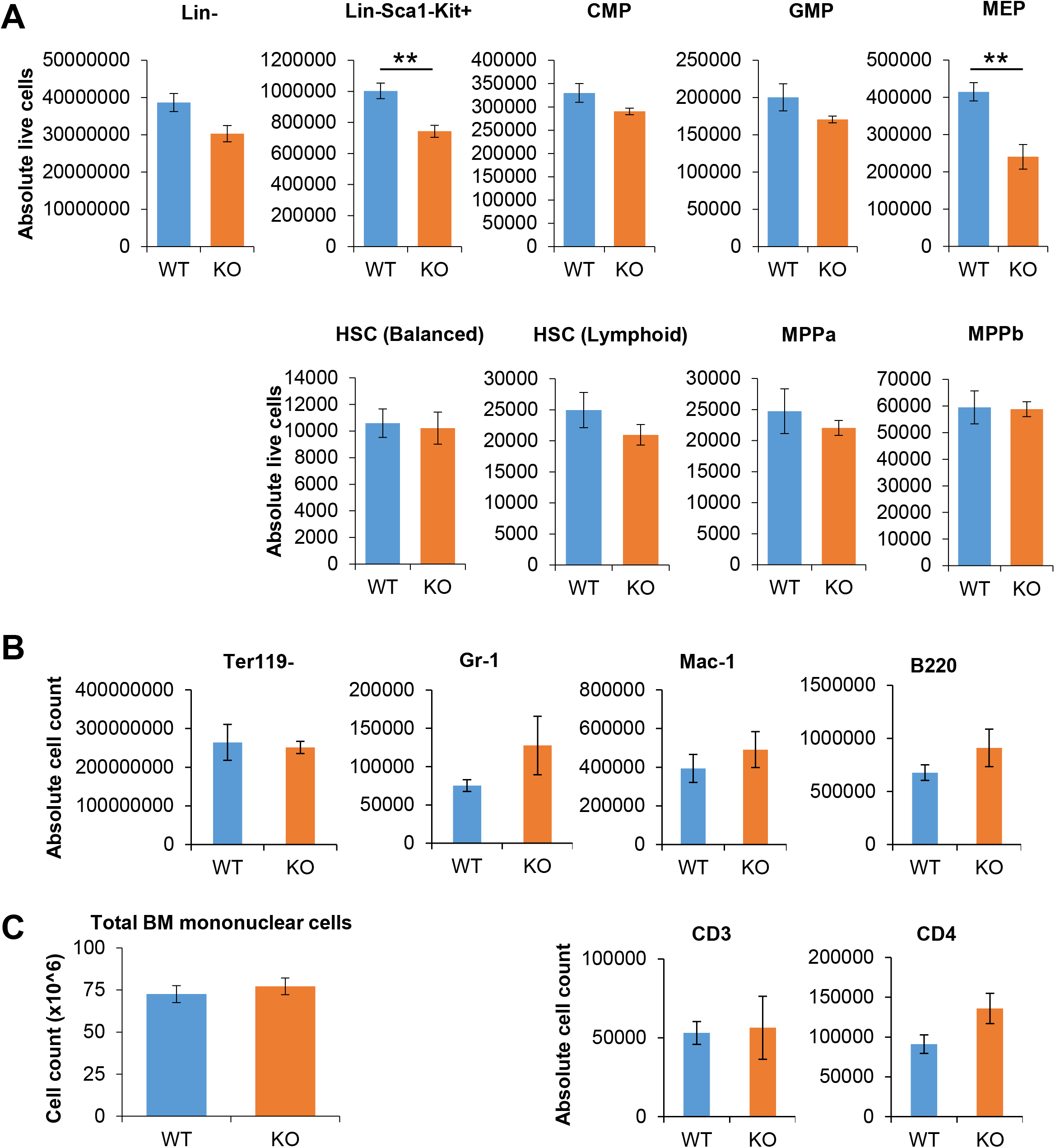

**Figure S3.**
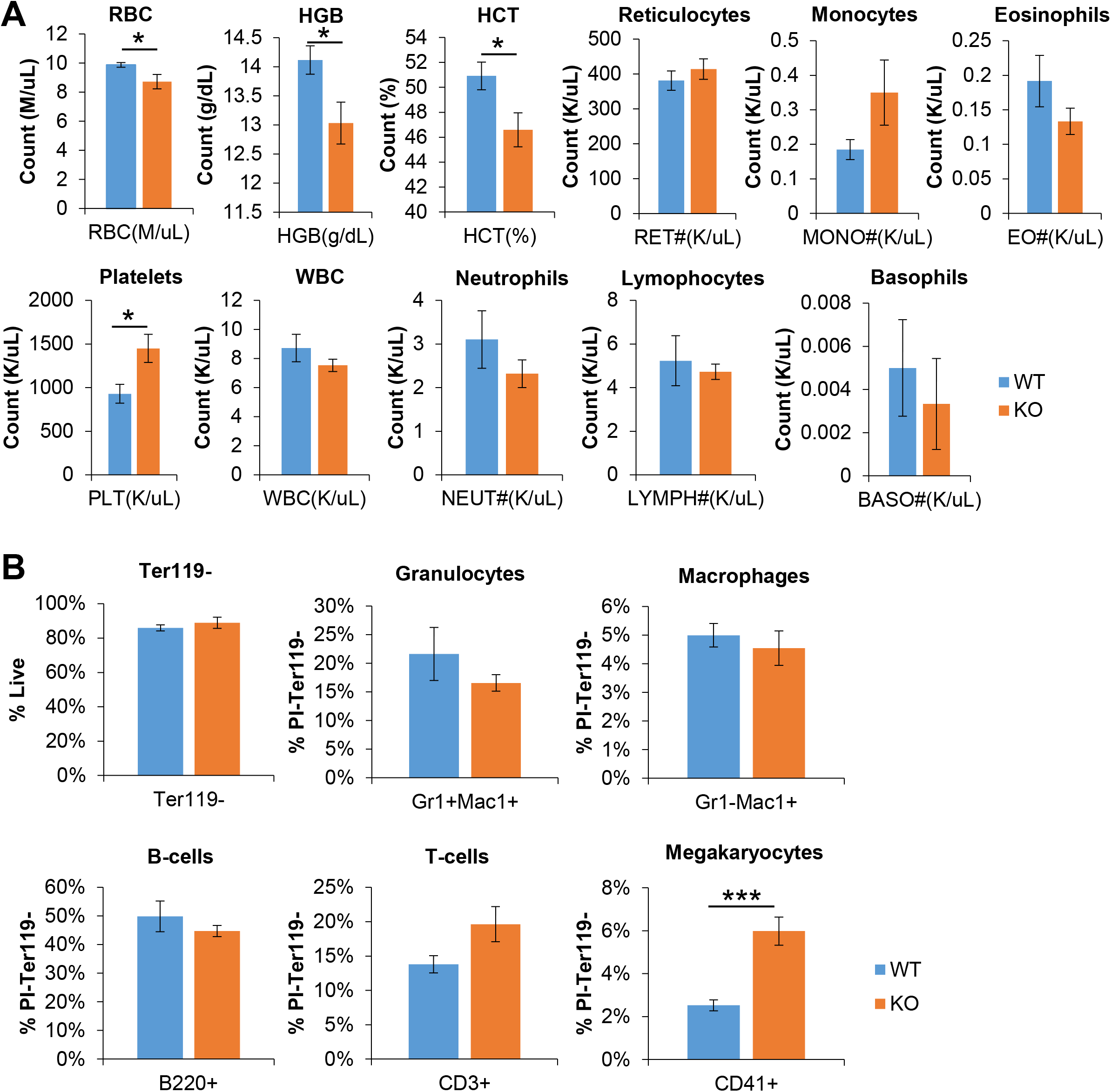

**Figure S4.**
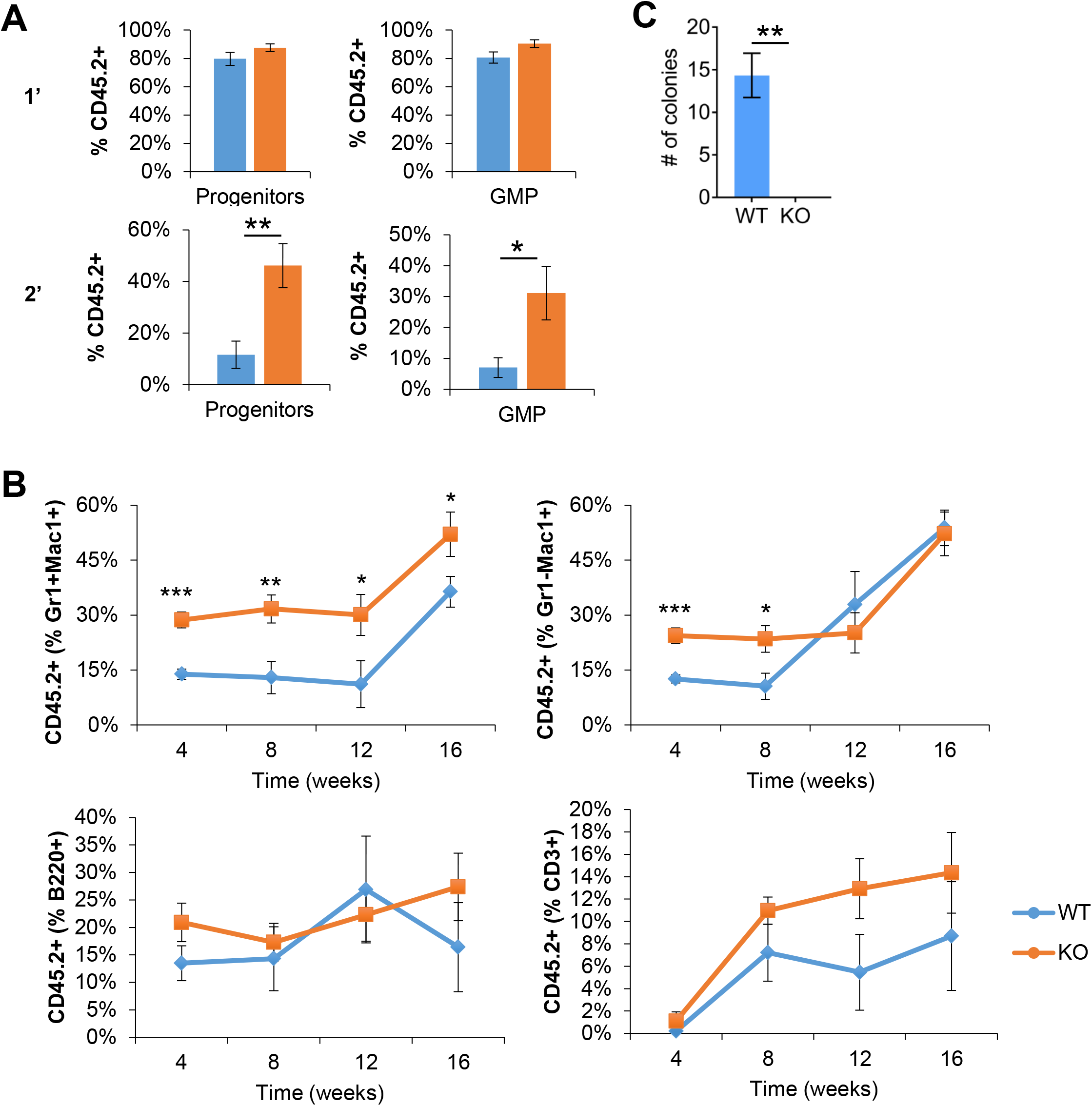

**Figure S5.**
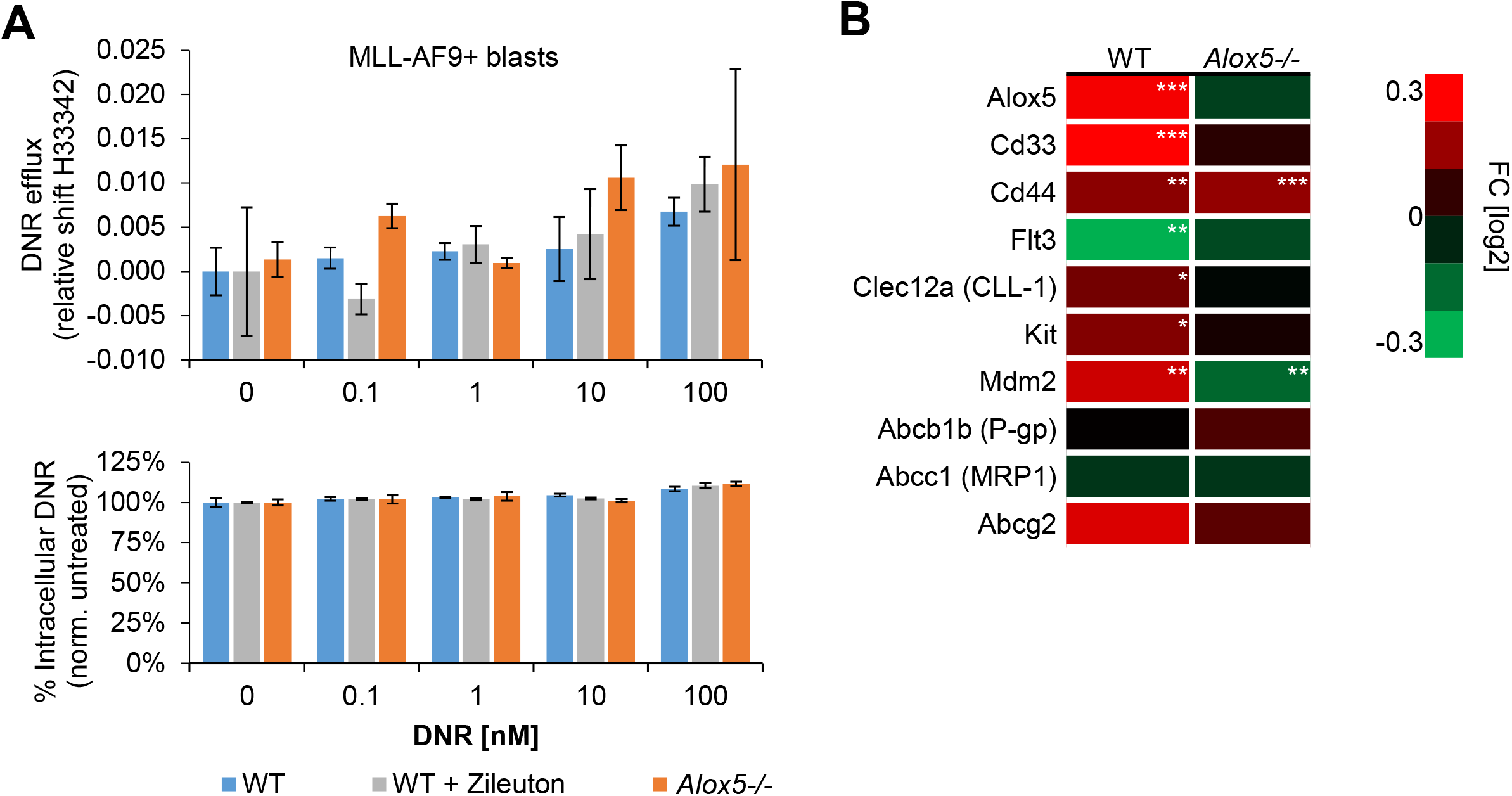

**Figure S6.**
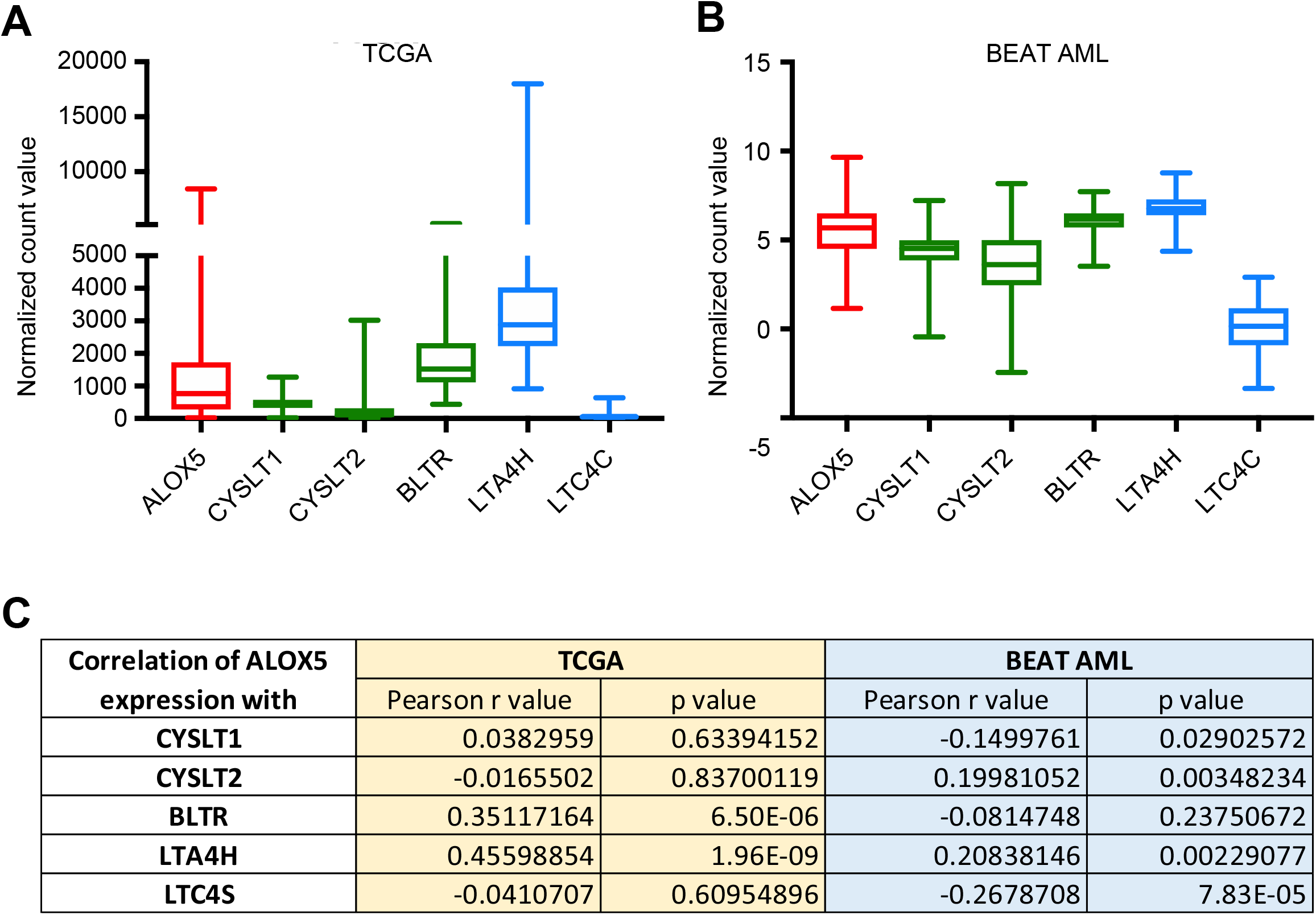

**Figure S7.**
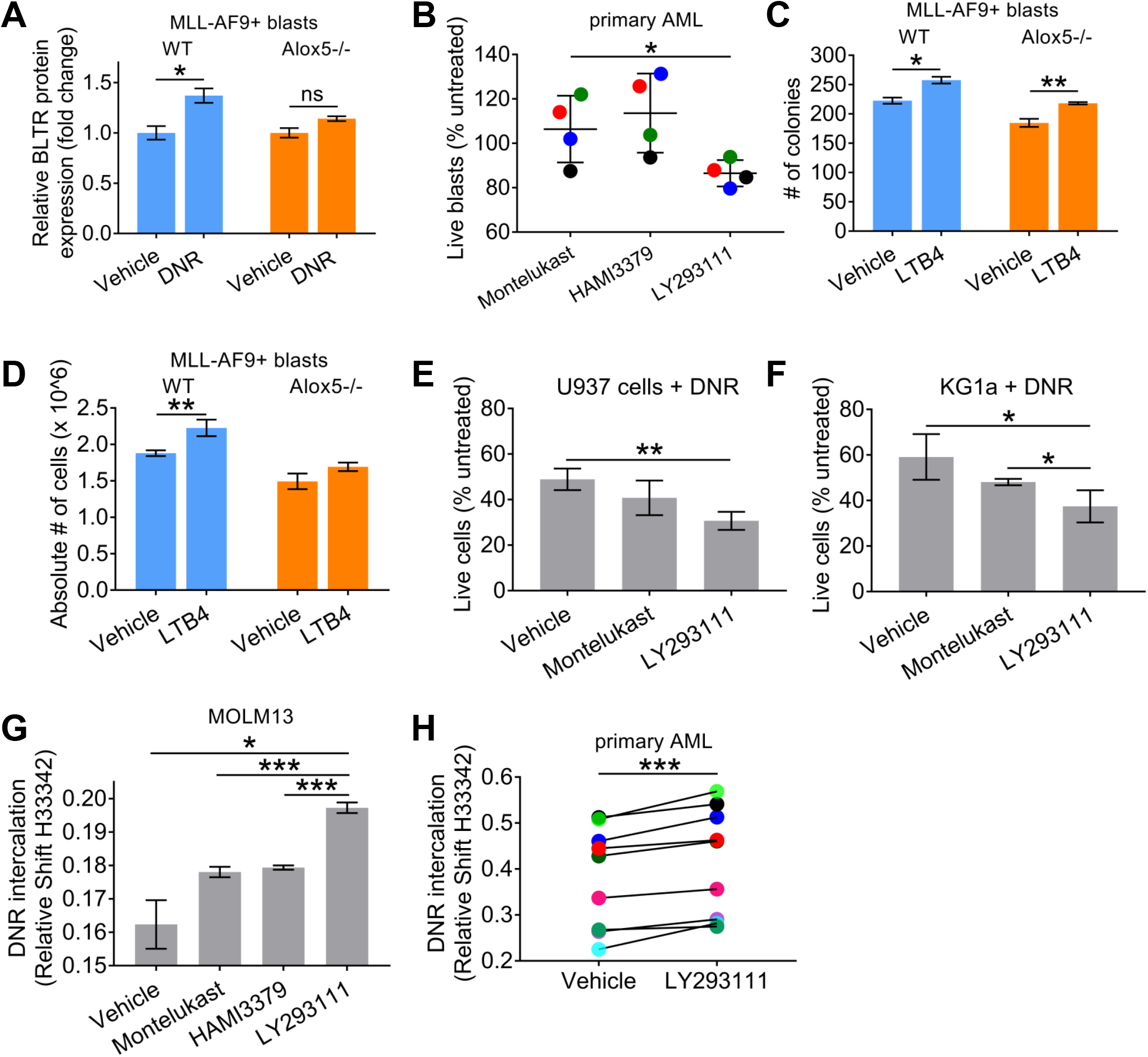

## References

1. Renneville A, Roumier C, Biggio V, Nibourel O, Boissel N, Fenaux P, et al. Cooperating gene mutations in acute myeloid leukemia: a review of the literature. Leukemia. 2008;22(5):915–31.

2. Bonnet D, Dick JE. Human acute myeloid leukemia is organized as a hierarchy that originates from a primitive hematopoietic cell. Nat Med. 1997;3(7):730–7.

3. Shlush LI, Mitchell A, Heisler L, Abelson S, Ng SWK, Trotman-Grant A, et al. Tracing the origins of relapse in acute myeloid leukaemia to stem cells. Nature. 2017;547(7661):104–8.

4. Pollyea DA, Jordan CT. Therapeutic targeting of acute myeloid leukemia stem cells. Blood. 2017;129(12):1627–35.

5. Kreitz J, Schonfeld C, Seibert M, Stolp V, Alshamleh I, Oellerich T, et al. Metabolic Plasticity of Acute Myeloid Leukemia. Cells. 2019;8(8).

6. Stefanko A, Thiede C, Ehninger G, Simons K, Grzybek M. Lipidomic approach for stratification of acute myeloid leukemia patients. PLoS One. 2017;12(2):e0168781.

7. Pabst T, Kortz L, Fiedler GM, Ceglarek U, Idle JR, Beyoglu D. The plasma lipidome in acute myeloid leukemia at diagnosis in relation to clinical disease features. BBA Clin. 2017;7:105–14.

8. Ye H, Adane B, Khan N, Sullivan T, Minhajuddin M, Gasparetto M, et al. Leukemic Stem Cells Evade Chemotherapy by Metabolic Adaptation to an Adipose Tissue Niche. Cell Stem Cell. 2016;19(1):23–37.

9. Stuani L, Riols F, Millard P, Sabatier M, Batut A, Saland E, et al. Stable Isotope Labeling Highlights Enhanced Fatty Acid and Lipid Metabolism in Human Acute Myeloid Leukemia. Int J Mol Sci. 2018;19(11).

10. Loew A, Kohnke T, Rehbeil E, Pietzner A, Weylandt KH. A Role for Lipid Mediators in Acute Myeloid Leukemia. Int J Mol Sci. 2019;20(10).

11. Hanna VS, Hafez EAA. Synopsis of arachidonic acid metabolism: A review. J Adv Res. 2018;11:23–32.

12. Chen Y, Hu Y, Zhang H, Peng C, Li S. Loss of the Alox5 gene impairs leukemia stem cells and prevents chronic myeloid leukemia. Nat Genet. 2009;41(7):783–92.

13. Yektaei-Karin E, Zovko A, Nilsson A, Nasman-Glaser B, Kanter L, Radmark O, et al. Modulation of leukotriene signaling inhibiting cell growth in chronic myeloid leukemia. Leuk Lymphoma. 2017;58(8):1903–13.

14. Zovko A, Yektaei-Karin E, Salamon D, Nilsson A, Wallvik J, Stenke L. Montelukast, a cysteinyl leukotriene receptor antagonist, inhibits the growth of chronic myeloid leukemia cells through apoptosis. Oncol Rep. 2018;40(2):902–8.

15. Wan M, Tang X, Stsiapanava A, Haeggstrom JZ. Biosynthesis of leukotriene B4. Semin Immunol. 2017;33:3–15.

16. Tager AM, Luster AD. BLT1 and BLT2: the leukotriene B(4) receptors. Prostaglandins Leukot Essent Fatty Acids. 2003;69(2-3):123–34.

17. Kanaoka Y, Boyce JA. Cysteinyl leukotrienes and their receptors: cellular distribution and function in immune and inflammatory responses. J Immunol. 2004;173(3):1503–10.

18. Austen KF. The cysteinyl leukotrienes: where do they come from? What are they? Where are they going? Nat Immunol. 2008;9(2):113–5.

19. Moore GY, Pidgeon GP. Cross-Talk between Cancer Cells and the Tumour Microenvironment: The Role of the 5-Lipoxygenase Pathway. Int J Mol Sci. 2017;18(2).

20. Haeggstrom JZ. Leukotriene biosynthetic enzymes as therapeutic targets. J Clin Invest. 2018;128(7):2680–90.

21. Massoumi R, Sjolander A. The role of leukotriene receptor signaling in inflammation and cancer. ScientificWorldJournal. 2007;7:1413–21.

22. Wang D, Dubois RN. Eicosanoids and cancer. Nat Rev Cancer. 2010;10(3):181–93.

23. Jala VR, Bodduluri SR, Satpathy SR, Chheda Z, Sharma RK, Haribabu B. The yin and yang of leukotriene B4 mediated inflammation in cancer. Semin Immunol. 2017;33:58–64.

24. Roos J, Oancea C, Heinssmann M, Khan D, Held H, Kahnt AS, et al. 5-Lipoxygenase is a candidate target for therapeutic management of stem cell-like cells in acute myeloid leukemia. Cancer Res. 2014;74(18):5244–55.

25. Rai KR, Holland JF, Glidewell OJ, Weinberg V, Brunner K, Obrecht JP, et al. Treatment of acute myelocytic leukemia: a study by cancer and leukemia group B. Blood. 1981;58(6):1203–12.

26. Prebet T, Bertoli S, Delaunay J, Pigneux A, Delabesse E, Mozziconacci MJ, et al. Anthracycline dose intensification improves molecular response and outcome of patients treated for core binding factor acute myeloid leukemia. Haematologica. 2014;99(10):e185–7.

27. Ishii S, Noguchi M, Miyano M, Matsumoto T, Noma M. Mutagenesis studies on the amino acid residues involved in the iron-binding and the activity of human 5-lipoxygenase. Biochem Biophys Res Commun. 1992;184(2):1133–4.

28. Nguyen T, Falgueyret JP, Abramovitz M, Riendeau D. Evaluation of the role of conserved His and Met residues among lipoxygenases by site-directed mutagenesis of recombinant human 5-lipoxygenase. J Biol Chem. 1991;266(32):22057–62.

29. Ziboh VA, Wong T, Wu MC, Yunis AA. Modulation of colony stimulating factor-induced murine myeloid colony formation by S-peptido-lipoxygenase products. Cancer Res. 1986;46(2):600–3.

30. Martin GH, Roy N, Chakraborty S, Desrichard A, Chung SS, Woolthuis CM, et al. CD97 is a critical regulator of acute myeloid leukemia stem cell function. J Exp Med. 2019.

31. Hu Y, Smyth GK. ELDA: extreme limiting dilution analysis for comparing depleted and enriched populations in stem cell and other assays. J Immunol Methods. 2009;347(1-2):70–8.

32. Agerstam H, Karlsson C, Hansen N, Sanden C, Askmyr M, von Palffy S, et al. Antibodies targeting human IL1RAP (IL1R3) show therapeutic effects in xenograft models of acute myeloid leukemia. Proc Natl Acad Sci U S A. 2015;112(34):10786–91.

33. Kikushige Y, Miyamoto T, Yuda J, Jabbarzadeh-Tabrizi S, Shima T, Takayanagi S, et al. A TIM-3/Gal-9 Autocrine Stimulatory Loop Drives Self-Renewal of Human Myeloid Leukemia Stem Cells and Leukemic Progression. Cell Stem Cell. 2015;17(3):341–52.

34. Majeti R, Chao MP, Alizadeh AA, Pang WW, Jaiswal S, Gibbs KD, Jr., et al. CD47 is an adverse prognostic factor and therapeutic antibody target on human acute myeloid leukemia stem cells. Cell. 2009;138(2):286–99.

35. Seita J, Sahoo D, Rossi DJ, Bhattacharya D, Serwold T, Inlay MA, et al. Gene Expression Commons: an open platform for absolute gene expression profiling. PLoS One. 2012;7(7):e40321.

36. Gentles AJ, Plevritis SK, Majeti R, Alizadeh AA. Association of a leukemic stem cell gene expression signature with clinical outcomes in acute myeloid leukemia. JAMA. 2010;304(24):2706–15.

37. Bagger FO, Kinalis S, Rapin N. BloodSpot: a database of healthy and malignant haematopoiesis updated with purified and single cell mRNA sequencing profiles. Nucleic Acids Res. 2019;47(D1):D881–D5.

38. Chung SS, Eng WS, Hu W, Khalaj M, Garrett-Bakelman FE, Tavakkoli M, et al. CD99 is a therapeutic target on disease stem cells in myeloid malignancies. Sci Transl Med. 2017;9(374).

39. Beerman I, Bhattacharya D, Zandi S, Sigvardsson M, Weissman IL, Bryder D, et al. Functionally distinct hematopoietic stem cells modulate hematopoietic lineage potential during aging by a mechanism of clonal expansion. Proc Natl Acad Sci U S A. 2010;107(12):5465–70.

40. Chen XS, Sheller JR, Johnson EN, Funk CD. Role of leukotrienes revealed by targeted disruption of the 5-lipoxygenase gene. Nature. 1994;372(6502):179–82.

41. Krivtsov AV, Twomey D, Feng Z, Stubbs MC, Wang Y, Faber J, et al. Transformation from committed progenitor to leukaemia stem cell initiated by MLL-AF9. Nature. 2006;442(7104):818–22.

42. Evans JF. Cysteinyl leukotriene receptors. Prostaglandins Other Lipid Mediat. 2002;68-69:587–97.

43. Haribabu B, Verghese MW, Steeber DA, Sellars DD, Bock CB, Snyderman R. Targeted disruption of the leukotriene B(4) receptor in mice reveals its role in inflammation and platelet-activating factor-induced anaphylaxis. J Exp Med. 2000;192(3):433–8.

44. Sofia MJ, Floreancig P, Bach N, Baker SR, Nelson K, Sawyer JS, et al. The discovery of LY293111, a novel, potent and orally active leukotriene B4 receptor antagonist of the biphenylphenol class. Adv Exp Med Biol. 1997;400A:381–6.

45. Jones TR, Labelle M, Belley M, Champion E, Charette L, Evans J, et al. Pharmacology of montelukast sodium (Singulair), a potent and selective leukotriene D4 receptor antagonist. Can J Physiol Pharmacol. 1995;73(2):191–201.

46. Wunder F, Tinel H, Kast R, Geerts A, Becker EM, Kolkhof P, et al. Pharmacological characterization of the first potent and selective antagonist at the cysteinyl leukotriene 2 (CysLT(2)) receptor. Br J Pharmacol. 2010;160(2):399–409.

47. Stenke L, Samuelsson J, Palmblad J, Dabrowski L, Reizenstein P, Lindgren JA. Elevated white blood cell synthesis of leukotriene C4 in chronic myelogenous leukaemia but not in polycythaemia vera. Br J Haematol. 1990;74(3):257–63.

48. Hashidate T, Murakami N, Nakagawa M, Ichikawa M, Kurokawa M, Shimizu T, et al. AML1 enhances the expression of leukotriene B4 type-1 receptor in leukocytes. FASEB J. 2010;24(9):3500-10.

49. Stenke L, Sjolinder M, Miale TD, Lindgren JA. Novel enzymatic abnormalities in AML and CML in blast crisis: elevated leucocyte leukotriene C4 synthase activity paralleled by deficient leukotriene biosynthesis from endogenous substrate. Br J Haematol. 1998;101(4):728–36.

50. Runarsson G, Feltenmark S, Forsell PK, Sjoberg J, Bjorkholm M, Claesson HE. The expression of cytosolic phospholipase A2 and biosynthesis of leukotriene B4 in acute myeloid leukemia cells. Eur J Haematol. 2007;79(6):468–76.

51. Dittmann KH, Mayer C, Rodemann HP, Petrides PE, Denzlinger C. MK-886, a leukotriene biosynthesis inhibitor, induces antiproliferative effects and apoptosis in HL-60 cells. Leuk Res. 1998;22(1):49–53.

52. Khan MA, Hoffbrand AV, Mehta A, Wright F, Tahami F, Wickremasinghe RG. MK 886, an antagonist of leukotriene generation, inhibits DNA synthesis in a subset of acute myeloid leukaemia cells. Leuk Res. 1993;17(9):759–62.

53. Wang Y, Skibbe JR, Hu C, Dong L, Ferchen K, Su R, et al. ALOX5 exhibits anti-tumor and drug-sensitizing effects in MLL-rearranged leukemia. Sci Rep. 2017;7(1):1853.

54. Sawyer JS, Bach NJ, Baker SR, Baldwin RF, Borromeo PS, Cockerham SL, et al. Synthetic and structure/activity studies on acid-substituted 2-arylphenols: discovery of 2-[2-propyl-3-[3-[2-ethyl-4-(4-fluorophenyl)-5-hydroxyphenoxy]-propoxy]phenoxy]benzoic acid, a high-affinity leukotriene B4 receptor antagonist. J Med Chem. 1995;38(22):4411–32.

55. Tong WG, Ding XZ, Hennig R, Witt RC, Standop J, Pour PM, et al. Leukotriene B4 receptor antagonist LY293111 inhibits proliferation and induces apoptosis in human pancreatic cancer cells. Clin Cancer Res. 2002;8(10):3232–42.

56. Hennig R, Ding XZ, Tong WG, Witt RC, Jovanovic BD, Adrian TE. Effect of LY293111 in combination with gemcitabine in colonic cancer. Cancer Lett. 2004;210(1):41–6.

57. Zhang W, McQueen T, Schober W, Rassidakis G, Andreeff M, Konopleva M. Leukotriene B4 receptor inhibitor LY293111 induces cell cycle arrest and apoptosis in human anaplastic large-cell lymphoma cells via JNK phosphorylation. Leukemia. 2005;19(11):1977–84.

58. Schwartz GK, Weitzman A, O’Reilly E, Brail L, de Alwis DP, Cleverly A, et al. Phase I and pharmacokinetic study of LY293111, an orally bioavailable LTB4 receptor antagonist, in patients with advanced solid tumors. J Clin Oncol. 2005;23(23):5365–73.

59. Pian P, Labovitz E, Hoffman K, Clavijo CF, Rzasa Lynn R, Galinkin JL, et al. Quantification of the 5-lipoxygenase inhibitor zileuton in human plasma using high performance liquid chromatography-tandem mass spectrometry. J Chromatogr B Analyt Technol Biomed Life Sci. 2013;937:79–83.

60. Winer ES, Stone RM. Novel therapy in Acute myeloid leukemia (AML): moving toward targeted approaches. Ther Adv Hematol. 2019;10:2040620719860645.

