## Supplemental Materials for "Leukotrienes promote stem cell self-renewal and chemoresistance in acute myeloid leukemia"

**Supplemental Methods**

**Human primary AML studies**

Human primary AML specimens were obtained from patients at Memorial Sloan Kettering Cancer Center, NYU Langone Health, and the University of Rochester (Rochester, NY) with informed consent under institutional approved protocols. Patients’ clinical characteristics, diagnosis, and molecular profiles are indicated in Supplemental Table 1. For survival studies, primary AML blasts were incubated in StemSpan media (StemSpan™ SFEM, cat no: 09600) containing cytokines (20 ng/ml hIL3, 20 ng/ml hIL6, 100 ng/ml hSCF, 50 ng/ml hTPO, 100 ng/ml hFLT3; Peprotech) for 72 hours.

**In vitro cell line studies**

Human cell lines Molm13, K562, U937, and KG1a were obtained from American Type Culture Collection (ATCC) or Deutsche Sammlung von Mikroorganismen und Zelkulturen GmbH (DSMZ) and maintained according to recommended guidelines. Cells were cultured in RPMI-1640 media (Thermo Scientific cat no: 11875119) containing 10% FBS (Sigma cat no: F2442) and antibiotics (penicillin 100 U/ml, streptomycin 100 U/ml, Thermo Scientific cat no: 10378016).

***Alox5*^-/-^ mouse studies**

*Alox5*^-/-^ mice (on C57BL/6J background) were purchased from The Jackson Laboratory (JAX, stock no: 004155). Homozygous deletion of Alox5 was confirmed by genotyping ear tissue by PCR. Mice were maintained in the Memorial Sloan Kettering Cancer Center animal facility per IACUC approved protocol.

*Alox5*^-/-^ serial transplantation studies were performed by transplanting double FACS-sorted HSCs (Lin^-^c-Kit^+^Sca-1^+^CD34^-^CD150^+^) isolated from WT or *Alox5*^-/-^ C57BL/6J mice. Lethally irradiated (2 x 475cGy = 950cGy total) PepBoyJ recipients were transplanted with 300 purified HSCs per mouse along with 2.4x10^5^ helper (PepBoyJ) total bone marrow cells by retro-orbital injection. Blood samples were collected every 4 weeks by tail vein bleed, and complete blood counts were assessed using an automated counter (Hemavet). Chimerism and lineage output were assessed by flow cytometry as described below. Mice were sacrificed 16 weeks following transplantation and secondary transplants were performed by injecting equal numbers of c-Kit^+^ cells from WT and KO recipients along with 2.5 x10^5^ helper (PepBoyJ) total bone marrow cells.

**Flow cytometry**

Cell sorting and analysis was performed using a BD FACSAria II sorter (BD Biosciences) and BD LSRFortessa (BD Biosciences), respectively. Antibody staining was performed for 60 minutes at 4°C in the dark. For cytotoxicity studies, propidium iodide (PI, 0.5µg/mL) and a fixed amount of unstained negative control beads (Beckman Coulter cat no: B22804) were added and a ratio of live cell to bead counts was measured and normalized to untreated controls (supplemental Table 3). Antibodies and other staining molecules used are shown in supplemental Table 4, and as described elsewhere(1). Flow cytometry data analysis was performed using FlowJo 9.8.5 software (FlowJo, LLC).

DNR DNA intercalation was measured as previously described(2, 3). Briefly, 1µM DNR was added to cultured cells with 5µM Hoechst 33342, resulting in quenching of the Hoechst 33342 signal based on the amount of DNR bound to DNA. Emissions were read in the indo-violet channel from the UV laser (355nm/20mW) and PE-Texas-Red channel from the Yellow-Green laser (561nm/50mW) for Hoechst and DNR, respectively, using 450 band pass (BP) and 575 BP filters, respectively. To quantitate the relative amount of DNR bound, transmission efficiency (TE) was calculated as TE = F0 – Fx/F0, where F0 and Fx are the mean fluorescence intensity (MFI) of the diploid peak for untreated and DNR treated samples, respectively. DNR efflux was calculated by incubating cells with DNR (0-1µM) for 60 minutes at 37°C, followed by a wash-out, and then incubating with Hoechst 33342 for 120 minutes at 37°C. Flow cytometric measurements were performed in the same manner as for the DNR binding studies.

**mRNA analysis**

ALOX5 transcript levels were quantified by qPCR. RNA was extracted from cells (Qiagen RNeasy Mini Kit, cat no: 74104) and cDNA was generated (ThermoFisher SuperScript III, cat no: 18080051) according to manufacturer’s specifications. qPCR was performed (ThermoScientific ABSolute Blue SYBR Green ROX Mix, cat no: AB-4162/B) with specific primers (supplemental Table 2). The ΔΔC_T_ method was used to calculate differences in gene expression with C_T_ measured by qPCR and expression of ALOX5 normalized to hPRT1. Normalized values were then compared to a reference sample and expression fold change was determined by converting from log_2_. Each reaction was performed in triplicate.

For RNA-sequencing analysis, WT MLL-AF9 and *Alox5*^-/-^ MLL-AF9 leukemic cells were treated *in vitro* with 5nM DNR for 72 hrs. RNA was extracted from cells (Qiagen RNeasy Mini Kit, cat no: 74104) and paired-end, 50 basepair RNA-sequencing was performed by Illumina^®^ Hiseq™ with SMARTer amplification at a read depth of 30-40x10^6^ reads per sample. Alignment metrics for each sample were calculated by GATK’s CollectRnaSeqMetrics and AlignmentSummaryMetrics. Sample clustering analysis was performed using multidimensional scaling of all samples. Heatmaps of the top 100 differentially-expressed genes in each comparison were generated for genes meeting fold change cutoff log_2_, adjusted p-value cutoff 0.05, and mean coverage of at least 15. Pathway analysis of all significantly changed genes (*P* < 0.05) was performed using the GSEA software package(4, 5). Differential gene expression comparisons were performed using the Gene List Venn Diagram software(6).

**Protein and leukotriene analysis**

Cells were centrifuged at 450 x g for 5 minutes at 4°C. Cell pellets were resuspended in 100ul NP-40 lysis buffer (10mM HEPES pH 7.9, 250mM NaCl, 5mM EDTA, 0.1% NP-40, 10% glycerol) and incubated on ice for 30 minutes. Lysates were spun at >1500 x g for 10 minutes at 4°C and supernatants were removed. Protein concentrations were quantitated by Bradford assay using BSA standard curves. Proteins were denatured in LDS buffer (C.B.S. Scientific ClearPage, cat no: FB31010) at 95°C for 5 minutes and loaded onto SDS-PAGE pre-cast gels (4-20% Mini-PROTEAN TGX Precast Protein Gel, cat no: 4561094). Proteins were transferred to 0.45µm nitrocellulose membrane, blocked with non-fat milk (BioRad) or 5% bovine serum albumin (Sigma), and washed with 0.1% Tween 20 Tris-buffered saline (pH 7.6). Primary and secondary antibodies used are listed in supplemental Table 4. ECL Western blotting detection reagents (Millipore) were used.

Leukotriene add-back experiments were performed in the presence of DNR (3 or 5nM) for 3 days with addition of LTB4 (10nM, Cayman Chemical cat no: 20110), LTC4 (100nM, Cayman Chemical cat no: 20210), LTD4 (100nM, Cayman Chemical cat no: 20310), and LTE4 (100nM, Cayman Chemical cat no: 20410) every 24 hours. Live cells were quantified by PI exclusion using flow cytometry.

**Supplemental Figures**

**Figure S1. Gene expression profiles of key enzyme encoding genes, in different AA pathways, in normal (CB, and BM), and leukemic (AML) cell types.** (A) ALOX5 for Lipoxygenase pathway, (B) CYP2J2 for Cytochrome P450/Epoxygenase pathway, and (C, D) PTGS1/2 for Cyclooxygenase pathway, as observed in Gene Expression Commons (<https://gexc.riken.jp/>).

**Figure S2. Alox5 KO mice exhibit normal bone marrow cellularity at steady-state.** (A) Absolute cell number of bone marrow stem and progenitor cell populations. (B) Absolute cell number of differentiated myeloid and lymphoid cell populations in the bone marrow of WT and *Alox5^-/-^* mice. (C) Total bone marrow mononuclear cell counts for WT and *Alox5^-/-^* mice. The results are presented as mean ± SEM. P-values were determined by the Student’s t-test: **P < 0.01.

**Figure S3. Alox5 KO mice show normal peripheral blood counts at steady-state.** (A) Complete blood counts of WT and *Alox5^-/-^* mice. (B) Frequency of myeloid and lymphoid cell populations in the peripheral blood of WT and *Alox5^-/-^* mice determined by flow cytometry. Results are presented as mean ± SEM. P-values were determined by the Student’s t-test: *P < 0.05; ***P < 0.001.

**Figure S4. Alox5 KO recipients exhibit elevated progenitor and myeloid lineage output in serial transplantation.** (A) Frequency of myeloid progenitors (Lin^-^Sca-1^-^c-Kit^+^) and GMP (Lin^-^Sca-1^-^c-Kit^+^CD34^+^CD16^+^) as a percentage of total CD45.2+ WT and *Alox5^-/-^* donor cells with serial transplantation. (B) Peripheral blood WT and *Alox5^-/-^* donor cell (CD45.2+ ) chimerism among granulocytes (Gr1^+^Mac1^+^), macrophages (Gr1^-^Mac1^+^) B-cells (B220^+^), and T-cells (CD3^+^). Tail bleeds were taken every 4 weeks for 16 weeks total. (C) Number of colonies as determined by CFU assay after plating WT or *Alox5*^-/-^ MLL-AF9 blasts. Primary (1’) and secondary (2’) transplantation. Results are presented as mean ± SEM. P-values were determined by the Student’s t-test: *P < 0.05, **P < 0.01, ***P < 0.001.

**Figure S5. Alox5 KO blasts exhibit a unique gene expression profile.** (A) Spectral shift of Hoechst 33342 peak emission with increasing concentration of DNR in WT and *Alox5^-/-^* MLL-AF9+ leukemic cells treated in vitro. (B) Evaluation of expression of known chemoresistance genes and canonical drug transporters, *P-*value only significant (*P*-value < 0.05) if indicated. The results are presented as mean ± SEM. P-values were determined by the Student’s t-test: *P < 0.05.

**Figure S6. ALOX5 transcript expression correlates with expression of genes from LTB4/BLTR axis in AML patient samples.** (A, B, C) Expression of ALOX5 and leukotriene receptor transcripts in AML patient samples as determined by TCGA and BEAT analysis.

**Figure S7. BLTR block sensitizes AML cell lines to the cytotoxic effects of DNR treatment**. (A) Normalized expression of BLTR protein in WT and *Alox5^-/-^* MLL-AF9+ leukemic blasts treated with vehicle or DNR *in vitro*. (B) Survival of patient AML blasts following treatment with Montelukast, HAMI3379, or LY293111. Number of colonies (C) and absolute number of cells (D) after plating WT or *Alox5-/-* MLL-AF9 blasts in Methocult containing either vehicle or 10 nM LTB4. (E,F) Survival of U937 and KG1a cells after treatment with DNR in combination with vehicle, CysLT1 receptor antagonist Montelukast or BLTR antagonist LY293111. (G) DNR DNA binding was measured following treatment of MOLM-13 cells with Montelukast, HAMI3379, or LY293111. (H) DNR DNA binding was assessed in primary AML patient blasts following treatment with LY293111. The results are presented as mean ± SEM. *P*-values were determined by the Student’s t-test: **P* < 0.05, ***P* < 0.01, ****P* < 0.001. Live cells were measured as % of PI- cells, normalized to untreated.

**Table S1. Patients’ clinical and molecular features.**

| **Fig.**  **S7B** | **Age** | **Cytogenetics** | **Gene** | **Variant**  **Effect** | **cDNA_Change** | **Protein_Change** | **FAF** |
| --- | --- | --- | --- | --- | --- | --- | --- |
| Pt. 1 | 56 | 46,XY,inv(16)(p13q22)[5]/47,idem,+14[16]nuc ish(ABL1,BCR)x2[200] | KIT | nonframeshiftBlock Substitution | c.1249_1254delACTTACinsCGGTCCTTTTTT | p.Thr417_Tyr418delinsArgSerPhePhe | 45.35 |
| Pt. 2 | 56 | 46,XX[20] | Not available | |  |  |  |
| Pt. 3 | 55 | 46,XY,inv(16)(p13q22)[18]/46,XX[2] nuc ish (CBFBx2)(CBFB5' sep CBFB3'x1)[154/200] | Not available | |  |  |  |
| Pt. 4 | 70 | 45,XY,-7[2]/49,XY,+6,+8,+9[2]/46,XY[17] | CSDE1,NRAS | missense, unknown | c.38G>A, c.*2046G>A | p.Gly13Asp, p.? | 16.56 |
|  |  |  | TET2 | nonsense | c.1852C>T | p.Gln618Ter | 18.5 |
|  |  |  | SRSF2 | missense, unknown | c.284C>G, c.-1476G>C | p.Pro95Arg, p.? | 48.86 |
|  |  |  | CEBPA | nonsense | c.625C>T | p.Gln209Ter | 45.67 |
| **Fig. 6F (CFU)** | | |  |  |  |  |  |
| Pt. 5 |  |  | PTPN11 | missense | c.226G>A | p.Glu76Lys | 41.37 |
| Pt. 6 | 74 | 46,XY,t(12;22)(p13;q11.2~12)[20] | NPM1 | Frameshift  Insertion | c.863_864insCCGG | p.Trp288CysfsTer12 | 42.6 |
|  |  |  | FLT3 | missense | c.2503G>C | p.Asp835His | 12.82 |
|  |  |  | IDH2 | missense | c.419G>A | p.Arg140Gln | 49 |
|  |  |  | SRSF2 | missense, unknown | c.284C>A, c.-1476G>T | p.Pro95His | 50.81 |
| Pt. 7 | 76 | 46,XX,der(7)t(1;7)(q25;q?31.2)[21] | NRAS | missense | c.297G>T | p.Gln99His | 49.55 |
|  |  |  | DNMT3A | missense | c.2645G>A | p.Arg882His | 49.55 |
|  |  |  | TET2 | missense | c.2599T>C | p.Tyr867His | 47.5 |
|  |  |  | TET2 | missense | c.5167C>T | p.Pro1723Ser | 51.28 |
|  |  |  | FLT3 | missense | c.2503G>T | p.Asp835Tyr | 45.92 |
|  |  |  | SRSF2 | missense, unknown | c.284C>A, c.-1476G>T | p.Pro95His, p.? | 52.15 |
|  |  |  | RUNX1 | missense | c.593A>G | p.Asp198Gly | 47.45 |
|  |  |  | BCOR | nonsense | c.3487C>T | p.Arg1163Ter | 47.52 |
|  |  |  | PHF6 | nonsense | c.820C>T | p.Arg274Ter | 43.72 |
| Pt. 8 |  | Not available |  |  |  |  |  |
| Pt. 9 | 74 | 46,XY,t(9;11)(p22;q23),?add(10)(q?24)[5]/46,XY[5] | PHF6 | frameshift | c.138delG | p.M46Ifs*35 | 96.12 |
| Pt. 10 |  | 46,XX[21] | NPM1 | INDEL |  | c.863_864insTCTG, p.W288CfsTer12 | 31.06 |
|  |  |  | FLT3 | FLT3-ITD 36(bp) |  | c.1800_1801dup, p.D600_L601insHFYVDFREYEYD | 2.9 |
|  |  |  | TET2 | INDEL |  | c.1403delA, p.H468LfsTer18 | 34.49 |
|  |  |  | TET2 | SNV |  | c.2728C>T, p.Q910Ter | 35.05 |
| **Fig. S7H (DNR binding)** | | |  |  |  |  |  |
| Pt. 1 | 56 | 46,XY,inv(16)(p13q22)[5]/47,idem,+14[16] nuc ish(ABL1,BCR)x2[200] | KIT | nonframeshiftBlock  Substitution | c.1249_1254delACTTACinsCGGTCCTTTTTT | p.Thr417_Tyr418delinsArgSerPhePhe | 45.35 |
| Pt. 2 | 56 | 46,XX[20] | Not available | |  |  |  |
| Pt. 3 | 55 | 46,XY,inv(16)(p13q22)[18]/46,XX[2] nuc ish (CBFBx2)(CBFB5' sep CBFB3'x1)[154/200] | Not available | |  |  |  |
| Pt. 4 | 70 | 45,XY,-7[2]/49,XY,+6,+8,+9[2]/46,XY[17] | CSDE1,NRAS | missense, unknown | c.38G>A, c.*2046G>A | p.Gly13Asp, p.? | 16.56 |
|  |  |  | TET2 | nonsense | c.1852C>T | p.Gln618Ter | 18.5 |
|  |  |  | SRSF2 | missense, unknown | c.284C>G, c.-1476G>C | p.Pro95Arg, p.? | 48.86 |
|  |  |  | CEBPA | nonsense | c.625C>T | p.Gln209Ter | 45.67 |
| Pt. 11 | 59 | 46,XY[20] |  |  |  |  |  |
| Pt. 12 | 66 | 46,XY,del(20)(q11.2q13.1)[20] | JAK2 | NON_SYNONYMOUS_CODING | c.1849G>T | p.V617F | 45.92 |
|  |  |  | PTPN11 | NON_SYNONYMOUS_CODING | c.179G>T | p.G60V | 38.91 |
| Pt. 13 | 68 | 45,X,-Y[5]/46,XY[3] | DNMT3A | missense_variant | c.2129G>A | p.C710Y | 43.06 |
|  |  |  | DNMT3A | stop_gained | c.1095C>G | p.Y365* | 45.2 |
|  |  |  | KIT | missense_variant | c.2447A>T | p.D816V | 34.85 |
|  |  |  | NPM1 | frameshift_variant | c.860_863dup | p.W288Cfs*12 | 36.57 |
|  |  |  | TET2 | missense_variant | c.2441G>A | p.R814H | 48.89 |
|  |  |  | TET2 | missense_variant | c.5704T>A | p.Y1902N | 41 |
|  |  |  | TET2 | frameshift_variant | c.4044delG | p.? | 43.24 |
| Pt. 14 | 67 | normal | FLT3 | inframe_insertion | c.1798_1799  insGGATGACCGGCTCCTCAGATAATGAGTACTTCTACGTTGATTTCAGAGAATATGAATATG | p.Y599_D600insGMTGSSDNEYFYVDFREYEY | 25.87 |
|  |  |  | IDH1 | missense_variant | c.394C>G | p.R132G | 48.76 |
|  |  |  | NPM1 | frameshift_variant | c.863_864 insCCTG | p.W288Cfs*12 | 46.81 |
| Pt. 15 | 79 | normal | U2AF1 | missense_variant | c.101C>T | p.S34F | 48.42 |
|  |  |  | FLT3 | disruptive_inframe_insertion | c.1743_1793  dupGACCGGCTCCTCAGATAATGAGTACTTCTACGTTGATTTCAGAGAATATGA | p.T582_E598dup | 28.79 |
|  |  |  | DNMT3A | missense_variant | c.2644C>T | p.R882C | 47.23 |
|  |  |  | ATM | missense_variant | c.7375C>T | p.R2459C | 51.26 |

| **Table S2. Primer sequences for Alox5 ectopic expression, mutagenesis, and qPCR** | | | |
| --- | --- | --- | --- |
| **Target** | **Primer** | **Sequence** | **Remarks** |
| ALOX5 | Forward | GCAGGAAGTGGCTACTGTGGA | Common (for genotyping) |
| ALOX5 | Forward | TGCAACCCAGTACTCATCAAG | Wild-type (for genotyping) |
| ALOX5 | Forward | ATCGCCTTCTTGACGAGTTC | Knockout (for genotyping) |
| ALOX5 | Forward | GTACGAATTCGTTTTCCCAGTCACGAC | Overexpression cloning |
| ALOX5 | Reverse | GCTAACCGGTCAGGAAACAGCTATGAC | Overexpression cloning |
| ALOX5 | Forward | GACCATCACCTCGCTTCTGCGAAC | Mutagenic PCR |
| ALOX5 | Reverse | GTTCGCAGAAGCGAGGTGATGGTC | Mutagenic PCR |
| pLentiLox |  | GCGATACTAGAGCTTGCATGC | Sequencing primer for ALOX5 incorporation |
| ALOX5 | Forward | ACTGGAAACACGGCAAAAAC | qPCR |
| ALOX5 | Reverse | TTTCTCAAAGTCGGCGAAGT | qPCR |
| hPRT1 | Forward | TCCAGCAGGTCAGCAAAGAA | qPCR |
| hPRT1 | Reverse | GAACGTCTTGCTCGAGATGT | qPCR |

| **Table S3. DNA Dyes for live/dead staining and DNR intercalation** | | | | |
| --- | --- | --- | --- | --- |
| **Dye** | **Conc.** | **Use** | **Source** | **Catalog No.** |
| DAPI | 0.01µg/mL | Cytotoxicity assay | Fisher Scientific | 26-829-810MG |
| PI | 0.5µg/mL | Cytotoxicity assay | Sigma-Aldrich | 81845 |
| H33342 | 5µM | Daunorubicin DNA binding assay | Sigma-Aldrich | B2261-100MG |

| **Table S4. Antibodies used for flow cytometry and western blot** | | | | | |
| --- | --- | --- | --- | --- | --- |
| **Antigen** | **Conjugate** | **Clone** | **Use** | **Source** | **Catalog No.** |
| c-Kit | PE/APC | 2B8 | Flow Cytometry | Biolegend | 105808/  105812 |
| Sca1 | PerCP | D7 | Flow Cytometry | Biolegend | 108122 |
| CD34 | APC | 581 | Flow Cytometry | Biolegend | 343510 |
| CD38 | PE | DL-101 | Flow Cytometry | Biolegend | 352306 |
| CD16/32 | A700 | 93 | Flow Cytometry | eBioscience | 56-0161-82 |
| Mac-1 | PB | M1/70 | Flow Cytometry | eBioscience | 48-0112-82 |
| Gr-1 | PE-Cy7/PE | RB6-8C5 | Flow Cytometry | Biolegend/  eBioscience | 108416/  12-5931-82 |
| B220 | PE-Cy5 | RA3-6B2 | Flow Cytometry | Biolegend | 103210 |
| CD3 | APC-Cy7 | 17A2 | Flow Cytometry | eBioscience | 47-0032-82 |
| Ter119 | PE-Cy7 | TER-119 | Flow Cytometry | Biolegend | 116222 |
| CD45 | A700 | HI30 | Flow Cytometry | Biolegend | 304024 |
| CD34 | APC-Cy7 | 561 | Flow Cytometry | Biolegend | 343614 |
| CD14 | BV421 | M5E2 | Flow Cytometry | Biolegend | 301830 |
| CD15 | PE-Cy7 | W6D3 | Flow Cytometry | Biolegend | 323030 |
| CD11b | APC | M1/70 | Flow Cytometry | Biolegend | 101212 |
| ALOX5 | - | Rabbit monoclonal | Primary in  Western Blot | Cell Signaling | 3289S |
| β-actin | - | Rabbit monoclonal | Primary in  Western Blot | Cell Signaling | 4970S |
| Rabbit IgG | HRP | Polyclonal | Secondary in  Western Blot | Cell Signaling | 7074P2 |
| Rabbit CysLT1 | - | Polyclonal | Flow Cytometry | Cayman | 120500 |
| Rabbit CysLT2 | - | Polyclonal | Flow Cytometry | Cayman | 120560 |
| Mouse BLTR | - | 202/7B1 | Flow Cytometry | Novus Biologicals | NB100-64831 |
| Goat anti-rabbit IgG | FITC | Polyclonal | Secondary in Flow Cytometry | ThermoFisher Scientific | 65-6111 |
| Goat anti-mouse IgG | A488 | Polyclonal | Secondary in Flow Cytometry | ThermoFisher Scientific | A11029 |

**Supplemental References**

1.	Shin JY, Hu W, Naramura M, Park CY. High c-Kit expression identifies hematopoietic stem cells with impaired self-renewal and megakaryocytic bias. J Exp Med. 2014;211(2):217-31.

2.	Belloc F, Lacombe F, Dumain P, Lopez F, Bernard P, Boisseau MR, et al. Intercalation of anthracyclines into living cell DNA analyzed by flow cytometry. Cytometry. 1992;13(8):880-5.

3.	Smeets ME, Raymakers RA, Vierwinden G, Pennings AH, Boezeman J, Minderman H, et al. Idarubicin DNA intercalation is reduced by MRP1 and not Pgp. Leukemia. 1999;13(9):1390-8.

4.	Subramanian A, Tamayo P, Mootha VK, Mukherjee S, Ebert BL, Gillette MA, et al. Gene set enrichment analysis: a knowledge-based approach for interpreting genome-wide expression profiles. Proc Natl Acad Sci U S A. 2005;102(43):15545-50.

5.	Mootha VK, Lindgren CM, Eriksson KF, Subramanian A, Sihag S, Lehar J, et al. PGC-1alpha-responsive genes involved in oxidative phosphorylation are coordinately downregulated in human diabetes. Nat Genet. 2003;34(3):267-73.

6.	Pirooznia M, Nagarajan V, Deng Y. GeneVenn - A web application for comparing gene lists using Venn diagrams. Bioinformation. 2007;1(10):420-2.
